# Basal cell of origin resolves neuroendocrine–tuft lineage plasticity in cancer

**DOI:** 10.1101/2024.11.13.623500

**Authors:** Abbie S. Ireland, Sarah B. Hawgood, Daniel A. Xie, Margaret W. Barbier, Scarlett Lucas-Randolph, Darren R. Tyson, Lisa Y. Zuo, Benjamin L. Witt, Ramaswamy Govindan, Afshin Dowlati, Justin C. Moser, Sonam Puri, Charles M. Rudin, Joseph M. Chan, Andrew Elliott, Trudy G. Oliver

## Abstract

Neuroendocrine and tuft cells are rare, chemosensory epithelial lineages defined by expression of ASCL1 and POU2F3 transcription factors, respectively^1,2^. Neuroendocrine cancers, including small cell lung cancer (SCLC), frequently display tuft-like subsets, a feature linked to poor patient outcomes^3–13^. The mechanisms driving neuroendocrine–tuft tumour heterogeneity, and the origins of tuft-like cancers are unknown. Using multiple genetically-engineered animal models of SCLC, we demonstrate that a basal cell of origin (but not the accepted neuroendocrine origin) generates neuroendocrine–tuft-like tumours that highly recapitulate human SCLC. Single-cell clonal analyses of basal-derived SCLC further uncovers unexpected transcriptional states and lineage trajectories underlying neuroendocrine–tuft plasticity. Uniquely in basal cells, introduction of genetic alterations enriched in human tuft-like SCLC, including high MYC, PTEN loss, and ASCL1 suppression, cooperate to promote tuft-like tumours. Transcriptomics of 944 human SCLCs reveal a basal-like subset and a tuft-ionocyte-like state that altogether demonstrate remarkable conservation between cancer states and normal basal cell injury response mechanisms^14–18^. Together, these data suggest that the basal cell is a plausible origin for SCLC and other neuroendocrine-tuft cancers that can explain neuroendocrine–tuft heterogeneity—offering new insights for targeting lineage plasticity.

## Main text

Tumour cell plasticity, an emerging cancer hallmark, promotes cell fate diversity, tumour progression, and resistance to therapy^19,20^. Precise cell-intrinsic and -extrinsic cues that dictate tumour cell plasticity remain unidentified for most cancers. If, and how, the cell-of-origin influences tumour plasticity is also largely uninvestigated.

Neuroendocrine cancers are typically aggressive tumours found in tissues throughout the body, including the lung, nose, breast, prostate, thymus, gastro-intestinal tract, pancreas and bladder^21–23^. Neuroendocrine tumours often exhibit a tuft-like subset of tumour cells that vary in proportion and are defined by expression of the lineage-related oncogene POU2F3^4–11,24–29^. How tuft-like cancers arise and the genetic alterations that may promote them are largely unknown, as there is currently a relative lack of model systems representing tuft-like cancer^30^. While diagnosis of tuft-like cancer is not yet clinically actionable, studies demonstrate that tuft-like cancers are transcriptionally distinct from their neuroendocrine counterparts, and preclinical studies suggest they have unique therapeutic vulnerabilities^9–11,31,32^.

Small cell lung cancer (SCLC) is one of deadliest neuroendocrine cancers, with a five-year survival rate of less than 7%^33–35^. SCLC comprises multiple molecular subtypes including ASCL1^+^ neuroendocrine, NEUROD1^+^ neuronal, and POU2F3^+^ tuft-like cell states (SCLC-A, -N, and -P, respectively)^34–37^. A fourth subset lacking ASCL1, NEUROD1, and POU2F3 is more controversial, and has been described as triple-negative, YAP1^+^, mesenchymal, and/or inflammatory^12,13,36,38–40^. Human SCLC exhibits intratumoural subtype heterogeneity, and several studies have demonstrated the capacity for tumour plasticity between SCLC-A, -N, and -Y states^12,41–50^, but whether neuroendocrine–tuft plasticity occurs is unknown.

An accepted cell of origin for SCLC is the pulmonary neuroendocrine cell (PNEC)^51–53^, a rare chemosensory population that expresses ASCL1^54,55^. However, introducing SCLC genetic alterations into PNECs is heretofore insufficient to generate SCLC-P in animal models^42,43^. SCLC-P is driven by tuft-lineage transcription factors^9,56,57^ and exhibits the worst survival among SCLC patients in response to both chemotherapy and chemo-immunotherapy^12,13,58^. SCLC-P is suspected to arise from rare chemosensory tuft/brush cells, so named for their apical microvilli that function in pathogen sensing and response^1^. But given the lack of animal models of tuft-like cancers, their cell of origin is currently unknown. Basal cells are the resident stem cell population of the lung that can differentiate during injury repair into common and rare cell types including ASCL1^+^ PNECs, POU2F3^+^ tuft cells, and FOXI1^+^ ionocytes^14–18,59–64^. Thus, homeostatic basal cell differentiation trajectories represent a largely unexplored connection linking neuroendocrine and tuft tumour fates. Whether a basal cell of origin enables neuroendocrine–tuft plasticity in SCLC is unknown.

### POU2F3^+^ tumours exhibit intratumoural subtype heterogeneity indicative of plasticity

Recent studies demonstrate that SCLC subtype evolution can occur between SCLC-A, -N, and -Y states^12,41–50^, but have failed to capture SCLC-P. To begin to determine if neuroendocrine–tuft plasticity occurs in SCLC, we assessed intratumoural heterogeneity of subtype-defining transcription factors in POU2F3^+^ human tumour biopsies. In a published analysis of 70 POU2F3-expressing human biopsies, >80% of tumours expressed at least one other SCLC subtype-defining transcription factor^50^. In our own hands, IHC analysis of 119 human SCLC biopsies revealed that ∼19% of tumours expressed POU2F3 (Extended Data Fig. 1a,b). Of the 23 POU2F3^+^ tumours, >82% co-expressed ASCL1 and/or NEUROD1 (Extended Data Fig. 1b). Co-immunofluorescence (co-IF) analysis on human biopsies (Extended Data Fig. 1c), in addition to POU2F3^+^ patient-derived xenografts (PDXs) (Extended Data Fig. 1d), revealed intratumoural heterogeneity of ASCL1 (A), NEUROD1 (N), and POU2F3 (P) that are often in mutually exclusive cell populations. Interestingly, rare double-positive A/N and A/P cells were detected in human biopsies and PDXs—indicative of potential transitional or intermediate cell states (Extended Data Fig. 1c,d). Thus, given the expected monoclonal origin of human tumours, intratumoural expression of POU2F3 with other subtype markers provides evidence that plasticity between SCLC-P and other subtypes likely occurs.

### Basal cells give rise to SCLC with expansive subtype heterogeneity

SCLC exhibits near universal loss of tumour suppressors *RB1* and *TP53,* as well as mutually-exclusive overexpression or amplification of a *MYC* family oncogene (commonly *MYCL* in neuroendocrine-high and *MYC* in neuroendocrine-low subsets)^65–67^. SCLC-P is associated with increased *MYC* amplification and overexpression^58^. Despite harbouring genetic drivers consistent with SCLC-P, MYC-high genetically-engineered mouse models (GEMMs) of SCLC (ie, *Rb1^fl/fl^/Trp53^fl/fl^/Myc^T58A^*(RPM) mice) demonstrate plasticity only between SCLC-A, -N, and -Y states in tumours arising from a neuroendocrine origin (using lineage-specific CGRP-Cre virus)^42,68^. POU2F3^+^ tumours are likewise undetectable in RPM GEMM tumours originating from alveolar or club cells^42,43,69^; however, rare POU2F3^+^ tumours arise in RPM GEMMs initiated by broadly active (non-lineage-specific) Ad-CMV-Cre^42^. Thus, an unknown cell of origin influences the tuft cell fate in SCLC.

During injury repair, basal cells give rise to both neuroendocrine and tuft lineages^14–18,59–64^. While neuroendocrine and tuft cells comprise <1% of the lung and tend to be post-mitotic^70–73^, basal cells represent ∼40% of lung cells in the major airways, ∼80% of proliferating lung cells, and are present throughout the human respiratory epithelium^14–17,59–63,74^, including at central airway branchpoints where SCLC is observed clinically^33,35^. Moreover, tobacco smoke, the primary risk factor for SCLC development^75,76^, increases the proliferative and inflammatory/metaplastic nature of lung basal cells^77^. Thus, we sought to determine whether basal cells can give rise to both neuroendocrine and tuft tumour cell fates.

While human basal cells are found throughout the major airways, mouse basal cells are located primarily in the tracheal epithelium^14,16,78^. To initiate SCLC from a basal cell in GEMMs, we employed the naphthalene injury model followed by basal-cell-specific adenoviral-*Krt5-*(K5)-Cre treatment, as described^79^ (Fig. 1a). In the RPM GEMM, tumours arose following K5-Cre with an average latency of 53 days, similar to the latency observed when tumours are initiated with CMV-(cell-type permissive), CGRP- (neuroendocrine-cell-specific), or CCSP- (club-cell-specific) Cre adenoviruses^43^ (Fig. 1b), with some RPM K5-Cre animals succumbing early to tumours occluding the trachea. K5-Cre initiated tumours formed in the trachea and major airways of the lung and exhibited SCLC histopathology (Fig. 1c).

**Fig. 1:**
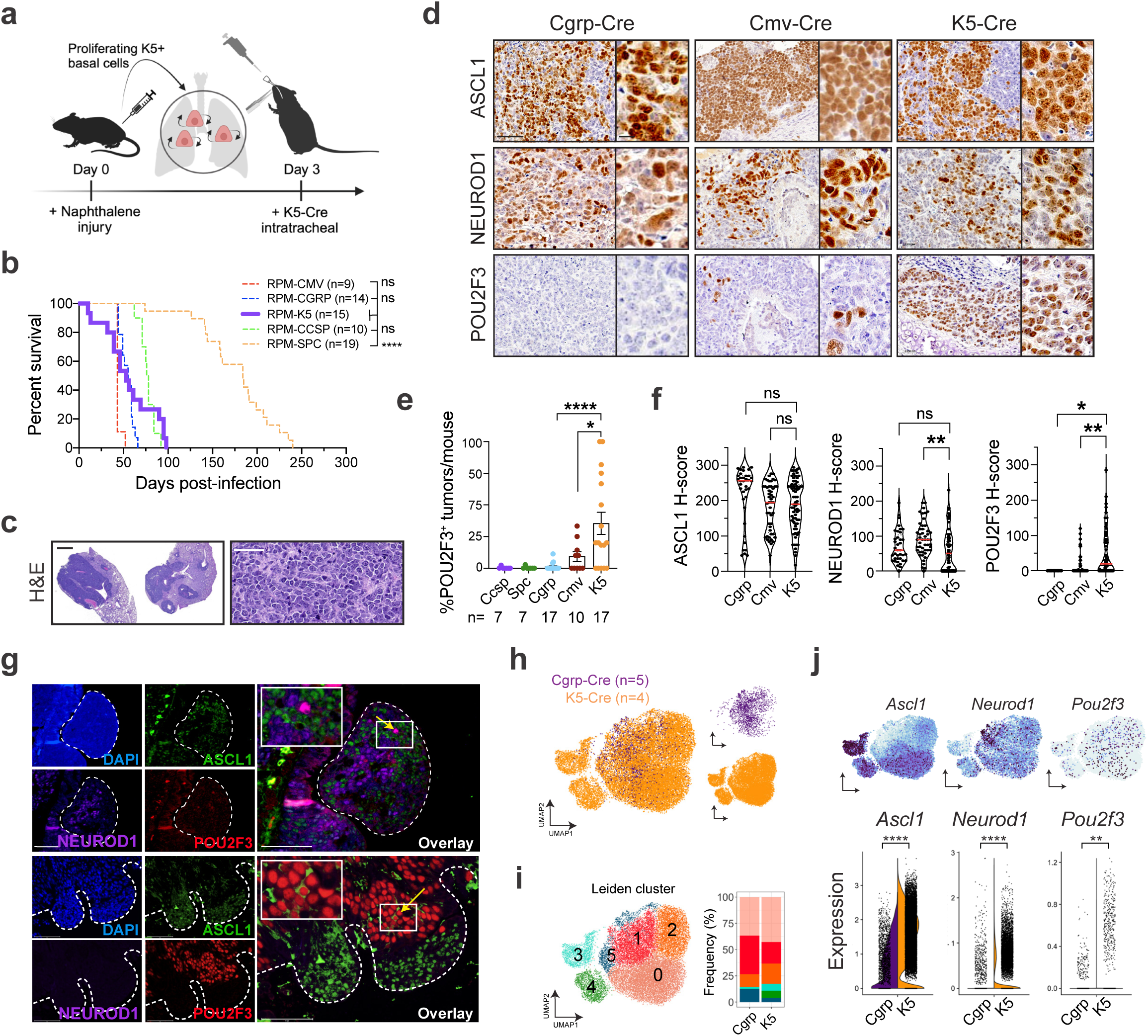
Basal cells give rise to SCLC with expansive subtype heterogeneity. See also Extended Data Fig. 2 and Supplementary Tables 1 and 2. **(a)** Schematic depicting method to induce SCLC from basal cells in the RPM GEMM. **(b)** Survival of RPM mice infected with indicated cell type-specific Ad-Cre viruses. Number of mice indicated in the figure. Dashed lines indicate historical data^43^. Mantel-Cox log-rank test comparing each cohort to K5-Cre (purple); **** p<0.0001; ns=not significant, p>0.05. **(c)** Hematoxylin and eosin (H&E) staining of RPM K5-Cre tumours. Representative whole lung lobes (left) and individual tumour morphology (right) depicted. Scale bars=1 mm (left) or 50 μm (right). **(d)** Representative IHC images from RPM tumours initiated by indicated Ad-Cre viruses for ASCL1, NEUROD1, and/or POU2F3. Scale bars=50 μm for main images, 10 μm for high magnification insets. **(e)** Quantification of number of POU2F3^+^ tumours (H-score >50) vs total tumour number per lung per mouse for indicated cell-type-specific or CMV-Cre adenoviruses. Each dot represents lungs of one mouse. Number of mice indicated in the figure. Error bars represent mean ± SEM. **(f)** H-score quantification of IHC on K5-Cre RPM tumours vs other cells of origin for indicated proteins. Each dot represents one tumour. For each marker, n=11-101 tumours quantified from n=4-19 mice per Ad-Cre group. Median (red bar) and upper and lower quartiles (dotted line) are indicated. **(g)** Representative co-immunofluorescent (co-IF) staining for DAPI (nuclei, blue), ASCL1 (green), NEUROD1 (purple), and POU2F3 (red) in RPM K5-Cre tumours. Tumour regions outlined with dashed line. High magnification insets of co-expressing cells (yellow arrows) are in the upper left corner of overlays. Scale bars=75 μm. **(h)** UMAP of scRNA-seq data from n=5 tumours initiated from NE cells (Cgrp-Cre, purple) from n=5 RPM mice, or n=4 tumours initiated from basal cells (K5-Cre, orange) from n=2 RPM mice. Cells coloured by sample in the same UMAP on right. **(i)** UMAP in (h) annotated by Leiden cluster (left) (Supplementary Table 1). Proportion of cells in Cgrp vs K5 tumours per Leiden cluster, represented as % of all cells per sample (right). **(j)** FeaturePlots depicting expression of indicated genes in UMAP, as in (h) (top). Split violin plot depicting mRNA expression of indicated genes by scRNA-seq for all cells in Cgrp (purple, left) and K5 (orange, right) tumours (bottom). Each dot is one cell. Student’s unpaired t-tests. **** p<0.0001, *** p<0.001, ** p<0.01, * p<0.05, ns=not significant, p>0.05. Unless otherwise noted, statistical tests are one-way ANOVA with Tukey’s correction (e,f). **** p<0.0001, ** p<0.006, * p<0.02, ns=not significant, p>0.05.

Immunostaining analysis of K5-initiated tumours revealed an absence of KRT5 (Extended Data Fig. 2a) and extensive heterogeneity in SCLC subtypes, including the presence of POU2F3^+^ tumours (Fig. 1d). Notably, POU2F3 was significantly more abundant in K5-initiated RPM tumours compared to other cells of origin (Fig. 1d,e).

We compared the abundance and intensity of A/N/P in RPM tumours from K5-versus CMV- or CGRP-Cre models and found significantly increased POU2F3 expression, slightly decreased but prominent expression of ASCL1 and NEUROD1, and less YAP1 (Fig. 1f and Extended Data Fig. 2b). Co-IF staining of RPM K5-Cre tumours revealed intratumoural heterogeneity of A, N, and P that predominantly mark distinct cells, with rare co-expressing cells (Fig. 1g), suggesting a potential transitory state. To assess transcriptional heterogeneity at the single cell level, multiple K5- and CGRP-initiated RPM tumours were profiled using single-cell RNA-seq (scRNA-seq) and projected via UMAP (Fig. 1h).

Leiden clustering of tumour cells suggests expanded transcriptional heterogeneity of tumours arising from basal versus neuroendocrine cells (Fig. 1h,i). Specifically, initiation from a basal cell enriched for cells in Leiden Clusters 3 and 4 (Fig. 1i). Leiden Cluster 3 highly expresses *Ascl1* but very little *Neurod1* whereas Cluster 4 has lower *Ascl1* levels but concomitant *Neurod1* expression (Fig. 1i,j). Compared to CGRP-initiated tumours, K5-initiated tumours exhibited increased expression of all SCLC subtype markers (Fig. 1j) and enrichment for SCLC-A and -P archetype signatures (derived from human SCLC cell lines)^46,80^, scRNA-seq-derived signatures of human SCLC-A and -P tumours^81^, and ASCL1 and POU2F3 ChIP-seq target genes^31,43,82^ (Extended Data Fig. 2c–e and Supplementary Table 2). K5-initiated tumours exhibited a slightly lower neuroendocrine score^83^ than CGRP-initiated tumours but remained predominantly neuroendocrine-high (Extended Data Fig. 2f). Together, these data support the hypothesis that the basal cell expands the subtype potential of RPM tumours, permitting intratumoural subtype heterogeneity of the major SCLC subtype states. Remarkably, basal-cell-initiated RPM tumours recapitulate both the spectrum of subtype heterogeneity and relative frequencies observed in human SCLC (A>N>P)—more so than any previously published SCLC GEMM.

### Basal-derived GEMM organoids give rise to neuroendocrine, neuronal, and tuft-like SCLC

To test the basal cell of origin more directly, we isolated normal tracheal basal cells from the RPM GEMM via surface ITGA6 expression^16,84^, an established basal cell marker (Fig. 2a), and plated cells in organoid culture conditions. Basal cells were transformed directly *ex vivo* with Ad5-CMV-Cre and subject to co-IF staining (Extended Data Fig. 3a–d) and scRNA-seq to determine phenotypic changes following transformation. Remarkably, transformed RPM basal organoids in culture exhibited few morphological or transcriptional differences compared to wildtype basal organoids (Fig. 2b and Extended Data Fig. 4a). Transformed cells had increased cycling, proliferation, and recombination markers (*Myc*, Firefly-luciferase-*fLuc*), but retained basal and stem markers, and low expression of other major lung cell type markers (AT1/2, club, goblet, ciliated) (Extended Data Fig. 4b,c and Supplementary Table 1).

**Fig. 2:**
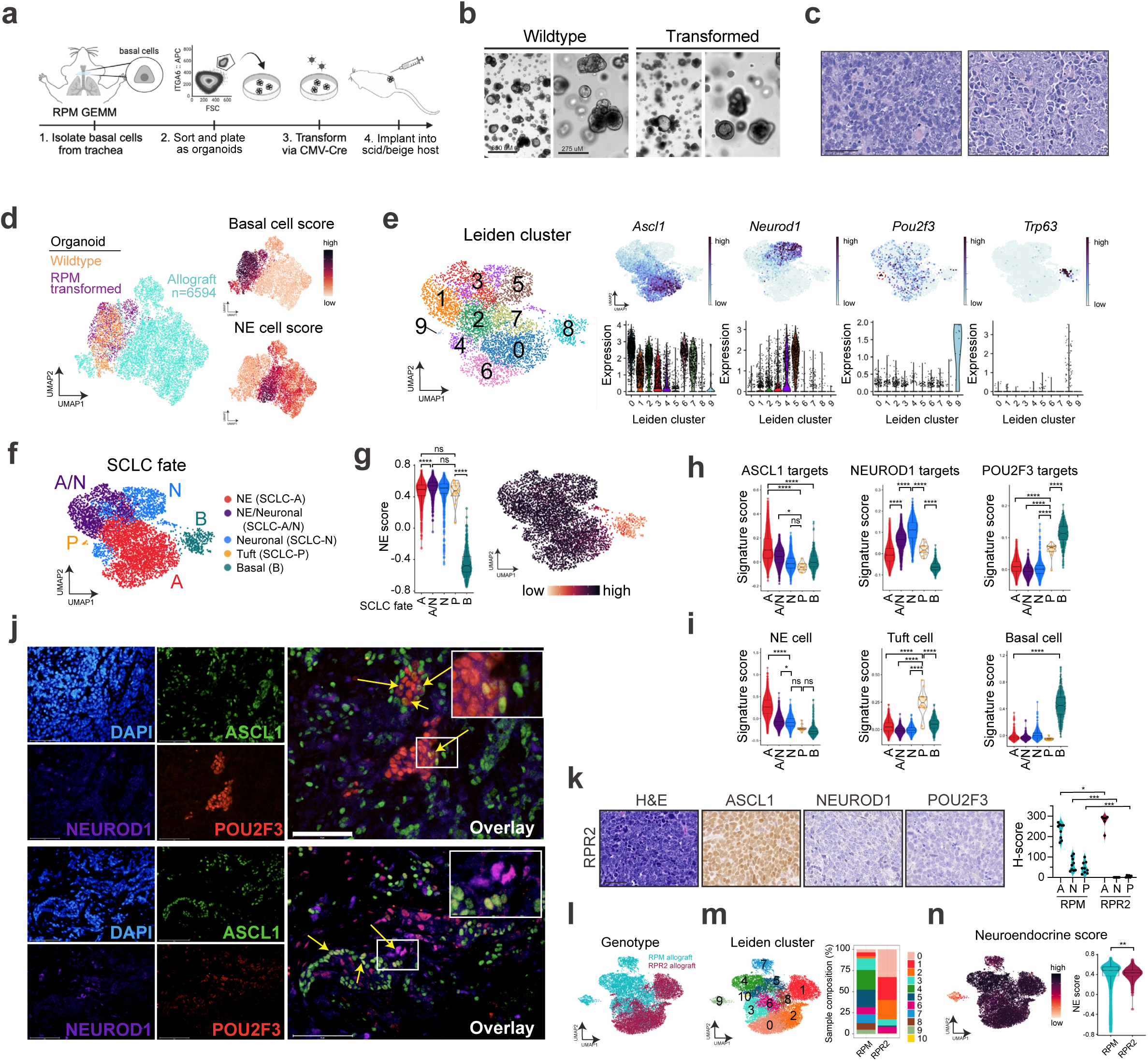
Basal-derived GEMM organoids give rise to neuroendocrine, neuronal, and tuft-like SCLC. See also Extended Data Figs. 3 and 4 and Supplementary Tables 1 and 2. **(a)** Schematic depicting isolation, growth and transformation of basal cell-derived organoids from RPM mice followed by implantation into the flanks of scid/beige hosts. **(b)** Representative brightfield images of basal organoids pre- (wildtype) and post- (transformed) CMV-Cre. Scale bars=650 μm (left, low mag) or 275 μm (right, high mag). **(c)** Representative H&E staining of RPM basal-organoid-derived tumours isolated from scid/beige mouse flanks with more classic (left) or variant (right) histopathology. Scale bar=50 μm. **(d)** UMAP of scRNA-seq data from wildtype (orange) and transformed (purple) RPM basal organoids, plus basal-organoid-derived RPM allograft tumour cells (turquoise). Allograft sample includes n=5 distinct RPM basal allograft tumours. FeaturePlots depicting expression of gene signatures derived from normal basal versus NE cells (right) (Supplementary Table 2). **(e)** UMAP of scRNA-seq data from RPM allograft tumours only, annotated by Leiden cluster (left) (Supplementary Table 1), FeaturePlot expression of indicated genes (top, right), and corresponding violin plot expression of indicated genes per Leiden cluster (bottom, right). Red dashed circle outlines Cluster 9 enriched for *Pou2f3*. **(f)** UMAP in (e) annotated by SCLC fate. Fates assigned based on enriched cell fate marker gene expression per Leiden cluster. **(g)** Violin plot of NE score^83^ per cell grouped by SCLC fate (left) from data in (f). UMAP of scRNA-seq data in (e) coloured by NE score (right). **(h)** Violin plot depicting ASCL1, NEUROD1, and POU2F3 ChIP target gene enrichment^31,42,43,113^ (Supplementary Table 2) in tumour cells from (e), grouped by SCLC fate assignment. **(i)** Violin plot depicting gene set enrichment of normal NE, tuft, and basal cells (Supplementary Table 2) in tumour cells from (e), grouped by SCLC fate assignment. **(j)** Representative co-immunofluorescent (co-IF) staining for DAPI (nuclei, blue), ASCL1 (green), NEUROD1 (purple), and POU2F3 (red) in RPM basal allograft tumours. High magnification insets of co-expressing cells (yellow arrows) are in the upper right corner of overlays. Scale bars=75 μm. **(k)** Representative IHC of RPR2 basal-derived allograft tumours for H&E and indicated SCLC subtype markers (left) with corresponding H-score quantification for ASCL1 (A), NEUROD1 (N) or POU2F3 (P) compared to RPM basal allograft tumours (right). Scale bar=50 μm. Mann-Whitney two-tailed t-test. * p<0.02, *** p<0.0005. **(l)** UMAP of scRNA-seq data from basal-organoid-derived RPM (turquoise, n=5 tumours) and RPR2 (maroon, n=1) allograft tumour cells. **(m)** UMAP in (l) annotated by Leiden cluster (Supplementary Table 1) (left). Proportion of cells from RPM vs RPR2 allograft tumours in each Leiden cluster, represented as % of all cells per sample (right). **(n)** UMAP of scRNA-seq data in (l) coloured by NE score (left). Violin plot of NE score per cell in RPM vs RPR2 basal allograft tumour cells (right). Student’s unpaired t-test. ** p<0.01. Box-whisker overlays on all violin plots indicate median and upper and lower quartile. Unless otherwise indicated, statistical tests are one-way ANOVA with Tukey’s correction. **** p<0.0001, * p<0.03, ns=not significant, p>0.05.

Consistent with transcriptomics, co-IF revealed significantly increased proliferation (KI67) in transformed RPM versus wildtype basal organoids, with maintained basal cell markers (DNP63, KRT8), and few to no cells expressing differentiated club (CCSP) or ciliated (FOXJ1) markers, respectively (Extended Data Fig. 3b,c). RPM wildtype and transformed basal organoids lacked SCLC-subtype markers by co-IF staining and scRNA-seq (Extended Data Figs. 3b–d and 4b), suggesting that transformed *Rb1/p53/Myc*-mutant cells maintain a basal fate under organoid conditions.

Transformed RPM basal organoids were implanted into the flanks of scid/beige mice to determine if they could generate SCLC tumours (Fig. 2a). Tumours arose in ∼4-6 weeks and histopathologically resembled SCLC (Fig. 2c). In contrast to organoids, basal-derived tumour allografts dramatically lose basal fate markers and gain neuroendocrine fate and proliferation markers (Fig. 2d and Extended Data Fig. 4d), suggesting environmental cues can dramatically impact tumour cell fate. To examine allograft heterogeneity at higher resolution, allograft tumour cells were reclustered without organoids (Fig. 2e).

The majority of Leiden clusters harboured cells expressing neuroendocrine (NE) and/or neuronal markers, while Cluster 9 cells expressed *Pou2f3* and tuft cell markers (Fig. 2e and Extended Data Fig. 4e). A small number of allograft cells in Cluster 8 also retained basal marker expression (Fig. 2e and Extended Data Fig. 4e). SCLC-A and -N archetype signatures were enriched in the majority of allograft clusters, while the -P archetype signature was enriched in Clusters 8 and 9 (Extended Data Fig. 4f). SCLC fates including SCLC-A, -N, -P, hybrid -A/N, and basal (B) were assigned per Leiden cluster (Fig. 2f)— informed by established markers and signatures of SCLC subtype states, normal lung cells, and differentially-expressed genes (Extended Data Fig. 4e,f and Supplementary Table 1). The *Ascl1^+^/Neurod1^+^* phenotype uniquely observed in RPM K5-Cre tumours (Fig. 1i, Cluster 4) was also represented in RPM basal allografts (Fig. 2f, “A/N”). This NE/Neuronal hybrid state has not been appreciated in mouse SCLC scRNA-seq data from other cells of origin, but is consistent with the frequent co-expression of ASCL1 and NEUROD1 in human tumours (Extended Data Fig. 1b), and with commonly observed *ASCL1^+^/NEUROD1^+^*human SCLC tumour cells by scRNA-seq^81^. In line with the assigned fates, application of the NE gene score revealed that most fates in RPM allograft tumours are NE-high, though cells in the tuft fate had a slightly lower NE-score, and basal state cells were NE-low (Fig. 2f,g).

Allografts exhibited variable ASCL1, NEUROD1, and POU2F3 ChIP-seq target gene signatures largely consistent with their assigned states (Fig. 2h). Allograft tumour fates were also appropriately enriched for signatures of their normal cell type counterparts^14^ (Fig. 2i). Characterization of RPM basal-derived allografts at the protein level revealed heterogeneous expression of SCLC subtype markers in largely mutually exclusive cell populations (Fig. 2j); similar to K5-Cre RPM primary tumours, RPM allografts predominantly expressed ASCL1 and NEUROD1, with minimal POU2F3 (Fig. 2j). Rare cells expressing a combination of A, N, and/or P (suggestive of intermediate or transitioning states) were again captured by co-IF (Fig. 2j). Together, both autochthonous and organoid-derived models suggest that neuroendocrine–tuft heterogeneity does not require a tuft-cell of origin and can derive from basal cells.

From neuroendocrine cells of origin, MYC can promote SCLC-A, -N, and -Y subtype plasticity in GEMMs, while non-MYC models such as *Rb1^fl/fl^/Trp53^fl/fl^/Rbl2^fl/fl^*(RPR2) are restricted to SCLC-A^42^. To test if MYC is required for SCLC subtype heterogeneity and the tuft fate specifically from a basal cell of origin, basal organoids were generated from RPR2 GEMMs that are known to express and occasionally amplify *Mycl*, but do not express MYC^42,68,85^. Transformed RPR2 organoids exhibit basal identity in vitro (Extended Data Figs. 3b–d, 4g), similar to RPM. Transplanted RPR2 allografts arose with significantly delayed latency (∼6 mo) compared to RPM allografts, consistent with their reduced latency in RPR2 autochthonous tumours^85^. In contrast to the subtype heterogeneity observed in RPM basal-derived allografts, RPR2 allografts were exclusively ASCL1^+^ with a relative lack of NEUROD1 and POU2F3 at the mRNA and protein levels (Fig. 2k and Extended Data Fig. 4h). RPR2 allografts also lacked *Myc* and expressed *Mycl*, in contrast to RPM allografts (Extended Data Fig. 4h). Leiden clustering of scRNA-seq data from RPR2 and RPM allografts revealed reduced transcriptional diversity in RPR2 tumours as they occupy fewer Leiden clusters (Figs. 2l,m and Supplementary Table 1) and lack cells in non-NE states compared to RPM allografts (Fig. 2n). Consistently, RPR2 allografts were enriched for the human SCLC-A archetype and had reduced SCLC-N and -P scores compared to RPM (Extended Data Fig. 4i).

Likewise, RPR2 allografts highly expressed ASCL1 ChIP target genes and had significantly reduced expression of POU2F3 and MYC target gene scores (Extended Data Fig. 4j). Altogether, these data suggest that MYC promotes tuft tumour fate specifically from a basal origin.

### ASCL1 loss promotes POU2F3^+^ tuft-like SCLC

ASCL1 is often mutually exclusive with the POU2F3^+^ tuft fate in cancer and development^4,6,14,24,48,58,81^ and is required for neuroendocrine phenotypes in multiple SCLC models^6,43,82,86,87^. Given that RPR2 and RPM basal allografts most commonly adopt the ASCL1^+^ NE-fate, we asked if ASCL1 inhibits tuft fate and whether its loss is sufficient to alter SCLC fate dynamics. Importantly, prior studies deleting *Ascl1* in RPR2 and RPM SCLC GEMMs in non-basal cells of origin did not enrich for POU2F3^43,82^.

To address the consequence of ASCL1 loss in basal-derived tumours, we generated organoids from *Rb1^fl/fl^/Trp53^fl/fl^/Myc^T58A^/Ascl1^fl/fl^* (RPMA) GEMMs^6,43^, transformed them with Ad5-CMV-Cre *ex vivo*, and established allografts as in Fig. 2a (Extended Data Figs. 3b-d and 5a). RPMA basal-derived allografts arose in 12-15 weeks, approximately twice the latency of RPM allografts (Extended Data Fig. 5b).

Resulting RPMA tumours exhibited predominantly SCLC histopathology (Fig. 3a). RPMA allografts lacked ASCL1 at the protein and mRNA levels, but intriguingly, still expressed NEUROD1 (Fig. 3b,c and Extended Data Fig. 5c); this is in stark contrast to findings from multiple non-basal cells of origin in the lung (i.e., NE, club, AT2), where loss of ASCL1 in RPM tumours completely blocks progression to the NEUROD1 state^43^. The presence of NEUROD1^+^ cells in RPMA basal allografts supports a capacity for the basal origin to alter SCLC fate potential compared to other cells of origin.

**Fig. 3:**
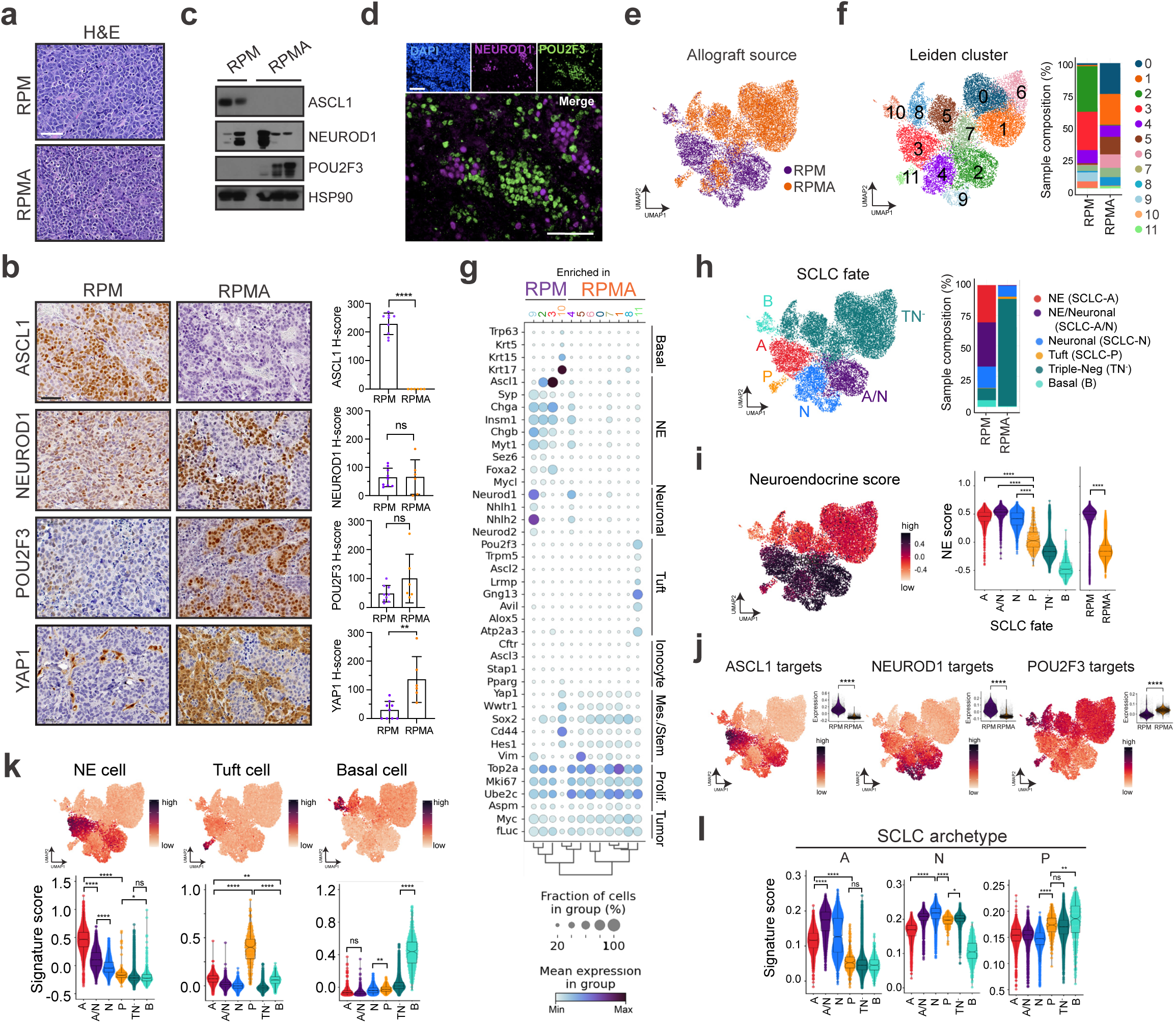
ASCL1 loss promotes POU2F3^+^ tuft-like SCLC. See also Extended Data Figs. 3 and 5 and Supplementary Tables 1 and 3. **(a)** Representative H&E staining of RPM (top) and RPMA (bottom) basal-organoid-derived tumours isolated from scid/beige mouse flanks. Scale bar=50 μm. **(b)** Representative IHC images from RPM and RPMA basal-organoid-derived tumours for indicated markers (left). H-score IHC quantification for indicated proteins (right). Each dot represents one tumour. For each marker, n=6-9 tumours quantified. Scale bars=50 μm. Student’s unpaired t-tests. **** p<0.0001, ** p<0.01, ns=not significant, p>0.05. Error bars represent mean ± SD. **(c)** Immunoblot depicting expression of indicated markers in RPM (n=2) vs RPMA (n=3) basal allograft tumours with HSP90 as a loading control. **(d)** Representative co-IF staining for DAPI (nuclei, blue), NEUROD1 (purple), and POU2F3 (green) in RPMA basal allograft tumours. Scale bars=75 μm. **(e)** UMAP of scRNA-seq data from basal-organoid-derived RPM (purple, n=5) and RPMA (orange, n=3) allograft tumours. **(f)** UMAP in (e) annotated by Leiden cluster (left). Proportion of cells from RPM vs RPMA allograft tumours in each Leiden cluster (Supplementary Table 3), represented as a % of all cells per sample (right). **(g)** Dot plot expression of genes marking indicated cell fates, stem-like, proliferative, and tumour cells for RPM and RPMA basal-derived allograft tumour cells, grouped by Leiden cluster as assigned in (f). Colour indicates level of gene expression and dot size represents frequency of expression per cluster. Genotypes indicate enrichment but not exclusive expression of each cluster. **(h)** UMAP of scRNA-seq data in (e) coloured by SCLC fate (left). Fates assigned based on enriched fate marker gene expression per Leiden cluster. Proportion of cells from RPM and RPMA allograft tumour samples in each fate, represented as % of all cells per sample (right). **(i)** UMAP of scRNA-seq data in (e) coloured by NE score according to legend (left). Violin plot of NE score per cell grouped by SCLC fate or genotype as indicated on the x-axis (right). **(j)** UMAP of scRNA-seq data in (e) coloured by ASCL1, NEUROD1, and POU2F3 ChIP target gene scores^31,42,43,113^ (Supplementary Table 2) where red/dark purple is high and orange is low. Upper right insets are violin plots depicting expression of target gene scores, grouped by genotype. Student’s unpaired t-tests. **** p<0.0001. **(k)** UMAPs (top) and violin plots (bottom) depicting gene set enrichment of normal NE, tuft, and basal cells (Supplementary Table 2) in tumour cells from (h), grouped by fate assignment. **(l)** Violin plot expression of SCLC subtype archetype signatures^46^ (Supplementary Table 2) per tumour cell in RPM vs RPMA basal allograft tumour samples from data in (h), grouped by SCLC fate. A=ASCL1, N=NEUROD1, and P=POU2F3. Box-whisker overlays on all violin plots indicate median and upper and lower quartile. Unless otherwise indicated, statistical tests are one-way ANOVA with Tukey’s correction. **** p<0.0001, ** p<0.001, * p<0.01, ns=not significant, p>0.05.

In contrast to basal-derived RPM allografts, deletion of ASCL1 promoted non-NE phenotypes of SCLC including robust and widespread expression of POU2F3 (Figs. 3b–d and Extended Data Fig. 5c)— emphasizing the importance of a basal origin for facilitating neuroendocrine–tuft heterogeneity. As in primary human tumours, expression of SCLC lineage-defining transcription factors was largely mutually exclusive within RPMA allografts (Fig. 3d). Other non-NE markers enriched in RPMA tumours included YAP1 (Fig. 3b), and Notch target gene product, HES1 (Extended Data Fig. 5d), which has a mutually antagonistic relationship with ASCL1 in lung development and neuroendocrine cancers^42,43,88–91^. Basal markers (KRT5, DNP63) were rarely observed in RPMA allografts, and were enriched, but not significantly altered compared to RPM tumours (Extended Data Fig. 5d).

ScRNA-seq and Leiden clustering on RPMA allograft cells compared to RPM shows depletion of clusters enriched for NE and neuronal cell markers and the emergence of a *Pou2f3^+^*state with global induction of key tuft-cell fate markers (Cluster 11; Fig. 3e–g and Extended Data Fig. 5e,f). Consistent with NEUROD1 expression at the protein level (Figs. 3b–d), one RPMA allograft cluster (Cluster 4) retained *Neurod1* and expression of other neuronal fate markers (Fig. 3e–g and Extended Data Fig. 5e,f). Leiden clusters were assigned to NE, Neuronal, hybrid NE/Neuronal, Tuft, or Basal fates, and the remaining RPMA-dominant clusters that were not clearly enriched for markers of these states were assigned to a Triple-Negative (TN^-^) state (Fig. 3h and Supplementary Table 3). This fate assignment and application of an NE score suggests that a majority of RPMA cells are in neuroendocrine-low *Pou2f3*^+^ (P) or subtype-negative (TN^-^) states, and a minority are in neuroendocrine-high Neuronal or hybrid NE/Neuronal states, which is highly divergent from RPM fate frequencies (Fig. 3h,i).

RPMA tumour cells exhibited a significant reduction of both ASCL1 and NEUROD1 ChIP target genes and a significant enrichment for POU2F3 target genes (Fig. 3j). Normal mouse cell type signatures and human SCLC archetype signatures were also enriched relatively consistent with assigned SCLC fate (Fig. 3k,l). Thus, loss of neuroendocrine fate driver ASCL1 is sufficient to shift the phenotypic landscape of SCLC towards a non-NE and SCLC-P state. While genetic alterations in *ASCL1* are not observed in human SCLC^65^, microenvironmental signals can repress neuroendocrine fate. For example, treatment with chemotherapy and alterations in Wnt^12,92^, Notch^42,88^, or YAP/TAZ^93–95^ signaling are known to repress ASCL1 and promote emergence of non-NE SCLC phenotypes, providing physiological relevance to the results from RPMA-derived tumours. Altogether, these data suggest that the basal cell of origin permits expansive SCLC heterogeneity reminiscent of differentiation paths taken by normal basal cells towards neuroendocrine and tuft cell fates. Prohibiting neuroendocrine differentiation of basal cells promotes emergence of SCLC-P, which to our knowledge represents the first *in vivo* SCLC model with a robust POU2F3^+^ population.

### Lineage-tracing reveals distinct SCLC evolutionary trajectories

Intratumoural heterogeneity and co-existence of mutually exclusive SCLC fates suggests lineage plasticity, but the mechanisms are relatively unknown. To determine if plasticity occurs between SCLC subtype states, we applied a lentiviral, combinatorial cell barcoding technology called CellTag^96,97^ to clonally trace basal-derived RPM and RPMA organoids and their resulting allografts at single-cell resolution (Fig. 4a,b). Organoids were infected with a multiplicity of infection >3 (yielding cells with a unique combination of three or more 8-bp CellTags), transformed with CMV-Cre, propagated to allow sufficient clonal expansion, then implanted into the flanks of scid/beige hosts (Fig. 4a). ScRNA-seq data from resulting RPM and RPMA organoids and allograft tumours (as depicted in Figs. 2–3) were subject to CellTag analysis. Fifty-five and 54 unique CellTagged clones were captured in RPM organoids and allografts, respectively, with 31 of the clones captured in both samples (Extended Data Fig. 6a and Supplementary Table 4). ScRNA-seq captured 63 unique clones in RPMA organoids and 89 in the resulting allograft, with 26 clones detected in both samples (Extended Data Fig. 6a and Supplementary Table 4). Since transformed organoids have minimal phenotypic diversity prior to transplantation (Extended Data Figs. 3–5), we focused on clones captured specifically in allograft tumours to interrogate fate plasticity.

**Fig. 4:**
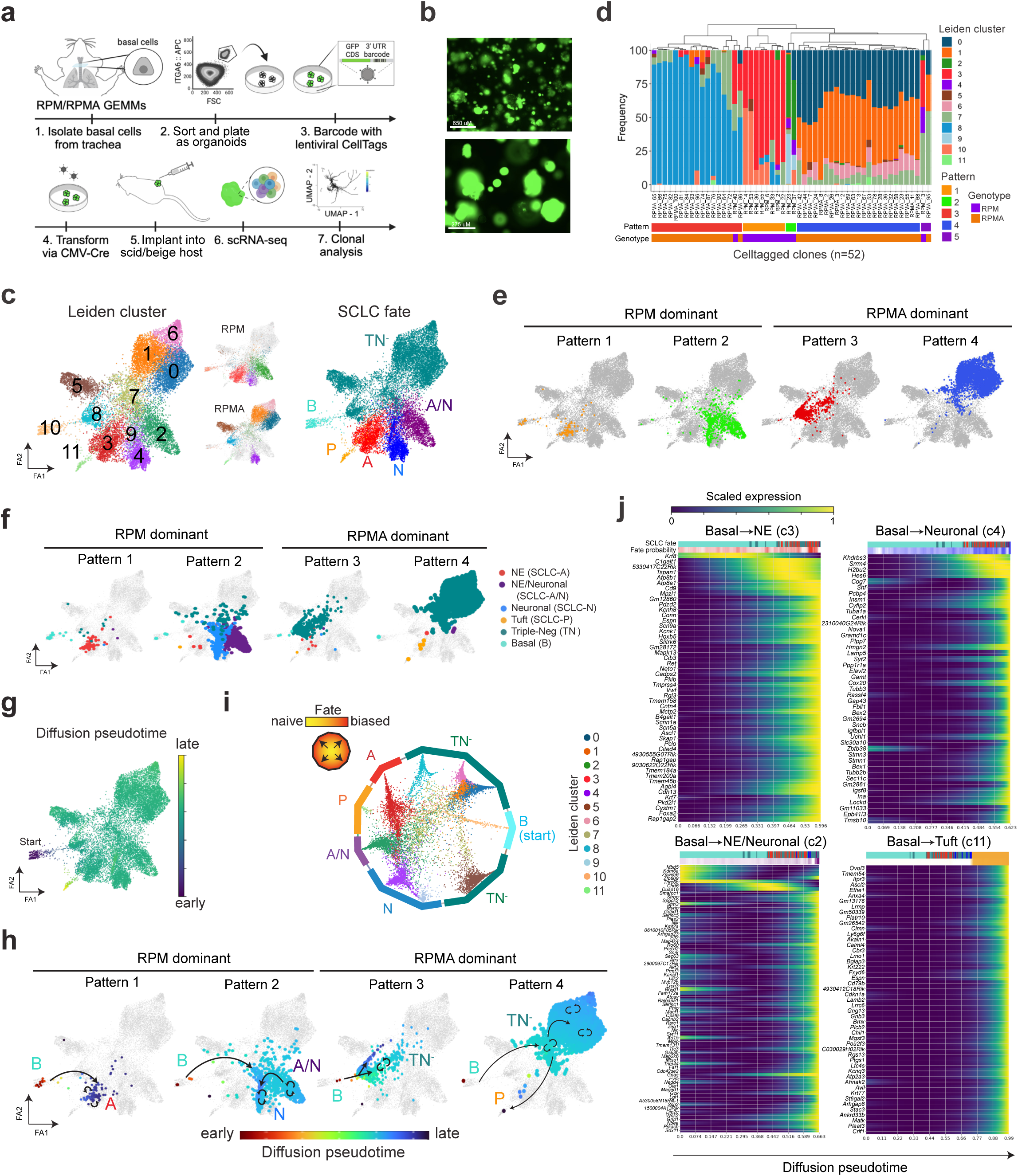
Lineage-tracing reveal distinct SCLC evolutionary trajectories. See also Extended Data Fig. 6 and Supplementary Tables 3–5. **(a)** Schematic depicting generation of CellTagged, basal RPM and RPMA organoids and allografts. **(b)** Representative fluorescent images of transformed and CellTagged RPM organoids (GFP coding sequence is included in the 3’ UTR of the CellTag library^96^). Scale bars=650 (top) and 275 (bottom) μm. **(c)** ForceAtlas2 (FA) map of RPM and RPMA basal allograft tumours annotated by Leiden cluster (as in Fig. 3f and Supplementary Table 3) (left) and split by genotype (middle). FA map of tumour cells coloured by assigned SCLC fate (as in Fig. 3h) (right). **(d)** Frequency of cells per Leiden cluster in each CellTag clone (one clone = one bar). Unbiased hierarchical clustering on Leiden cluster occupancy of all clones reveals four major patterns (Pattern 1-4) and one minor pattern (Pattern 5), labeled on the x-axis. Genotype and unique number of each clone indicated on x-axis and matching clone numbers in Extended Data Fig. 6b,c. **(e)** FA maps of major clonal Patterns according to assignment and colour in (d). Patterns dominated by RPM vs RPMA-derived clones are labeled. **(f)** FA maps as in (c) of RPM and RPMA basal allograft clonal dynamics grouped by clonal dynamic Pattern and annotated by corresponding SCLC fate. **(g)** FA map as in (c) coloured by diffusion pseudotime, implemented in Scanpy. Assigned pseudotime starting state was basal-enriched Cluster 10. **(h)** Clonal dynamic Patterns as in (e) but annotated by pseudotime assignment from (g). Arrows drawn on plots follow predicted pseudotime trajectories, with straight arrows indicating movement of cells across Leiden clusters and circular arrows indicating self-renewal/proliferation within that cell state or Leiden cluster. Fate assignments also annotated as in (c). **(i)** CellRank analysis of fate probabilities per cell in scRNA-seq data of RPM and RPMA basal allograft tumours from (c). Each dot represents a tumour cell, coloured by Leiden cluster. Cells are arranged inside the circle according to fate probability, with fate-biased cells placed next to their corresponding edge and naive cells located in the middle. Edges annotated by SCLC fate as in (c). **(j)** CellRank trajectory-specific expression trends of putative driver genes (Supplementary Table 5), predicted by fitting gene expression and pseudotime coordinates with Generalized Additive Models (GAMs).

RPM and RPMA allograft tumour cell transcriptomic data were projected via ForceAtlas2 (FA) force-directed graphing^98^ (Fig. 4c)—an alternative to UMAP that preserves data topology and is often preferred for trajectory analyses. To interrogate clonal dynamics, we visualized the frequency of cells per Leiden cluster in each individual RPM and RPMA CellTagged clone (defined as >5 cells per clone post-QC; n=12 RPM, n=40 RPMA) (Fig. 4d and Extended Data Fig. 6b,c). Unbiased clustering of clones by Leiden occupancy revealed four major patterns (Pattern 1-4) and one minor pattern (Pattern 5) (Fig. 4c–e)—suggesting cells evolve through defined plasticity patterns rather than randomly through transcriptional space.

To assess how the clonal dynamics of each pattern correspond with fate plasticity, SCLC fates were annotated in FA-embedded space and visualized per clonal dynamic pattern (Fig. 4c,f and Extended Data Fig. 6d). Pattern 1 and 2 clones were exclusively from RPM tumours, while Pattern 3 and 4 clones largely represented RPMA clonal dynamics (Figs. 4c–f). Pattern 1 clones largely exhibited plasticity between Basal and *Ascl1*^+^ NE fates, corresponding with Leiden clusters 10 and 3, respectively (Fig. 4c,f). In contrast to Pattern 1, Pattern 2 RPM clones exhibited expanded plasticity between Basal, hybrid NE/Neuronal, and Neuronal states, with one cell in the Tuft state and a few in the TN^-^ state (Fig. 4c,f and Extended Data Fig. 6b)—suggesting that clones with the majority of cells in the SCLC-A state have limited fate potential compared to clones with cells in the hybrid SCLC-A/N state. Together, the clonal dynamics of RPM cells suggest that basal cells have a propensity to adopt an SCLC-A state or to adopt a hybrid NE/Neuronal phenotype that can also access Neuronal or non-NE (P or TN^-^) fates.

RPMA clones exhibit distinct transcriptional plasticity patterns compared to RPM (Fig. 4c–e). Pattern 3 clones were primarily TN^-^, with very few cells in the NE and NE/Neuronal clusters, consistent with loss of *Ascl1* (Fig. 4f). Pattern 4 clones also had an abundance of cells in the TN^-^ state, which was the most common clonal pattern to give rise to Tuft state cells (Fig. 4f and Extended Data Fig. 6c). Taken together, RPMA clones exhibit greater plasticity between TN^-^ and Tuft states than RPM clones.

The CellTag-based clonal analysis as performed has limited insights into the *directionality* of subtype switching, but predicted directionality of plasticity was determined using diffusion pseudotime analysis (Fig. 4g). From an assigned starting state of Basal identity (Cluster 10), pseudotime predicts Pattern 1 plasticity from Basal to NE, Pattern 2 from Basal to hybrid NE/Neuronal to Neuronal, Pattern 3 from Basal to TN^-^ (Cluster 7 then Cluster 8), and Pattern 4 from Basal to TN^-^ (Cluster 7 then Clusters 0, 1, 6) to Tuft (Cluster 11)(Fig. 4h).

Lastly, to infer the probability of tumour cells in RPM and RPMA allografts reaching each assigned fate, and to identify fate drivers, we applied CellRank analysis^99^. CellRank predicted varying levels of fate determination along multiple trajectories starting in the basal state (Extended Data Fig. 6e). Many tumour cells were heavily fate biased, but some remained fate naïve (or undifferentiated/uncommitted) (Fig. 4i). Cells in Neuronal, Tuft, and NE clusters were particularly biased towards their respective fates (closer to their respective edges) (Fig. 4i). In contrast, Basal, TN^-^, and hybrid NE/Neuronal clusters had both fate-biased and -naïve cells. Cluster 7 TN^-^ cells were the most unbiased (most central), suggesting Cluster 7 cells represent a lineage-naïve state (Fig. 4i). Interestingly, the majority of CellTagged clones had some cells in Cluster 7 (Fig. 4c,d), and diffusion pseudotime analyses predict cells adopt this state prior to reaching more fate-biased states (Fig. 4h). Analysis of marker genes for Cluster 7 demonstrate enrichment of metabolic, neuronal progenitor, and cycling basal cell signatures (Supplementary Table 3). Notably, this cluster appears to be a “lineage-naïve” or “lineage-confused” state at the transcriptional level; whether this state is related to the “highly-plastic states” observed in other cancer types^100^ warrants further study.

CellRank fits gene expression data of cells ordered in pseudotime to Generalized Additive Models (GAMs)^101^ to identify and compute expression of putative driver genes along predicted differentiation trajectories. Predicted NE lineage drivers include early induction of *Krt8* followed by gain of *Ascl1*, *Vwf*, and *Foxa2* (Fig. 4j); supporting this prediction, FOXA2 is present in ASCL1-high SCLC super enhancers^82^ and can interact with ASCL1 to regulate NE programs in prostate neuroendocrine cancer^102^. Neuronal fate drivers include *Srrm4* (a neuronal-specific splicing factor implicated in REST splicing^103^), *Hes6*, *Tubb3*, and *Uchl1* (Fig. 4j); consistently, SRRM4 expression highly correlates with a NEUROD1^+^ subset of prostate neuroendocrine cancer^104^. Interestingly, drivers of differentiation towards a hybrid NE/Neuronal state include early activation of *Kdm6a*, followed by induction of *Kmt2a* (Fig. 4j), both recently implicated in ASCL1-to-NEUROD1 lineage plasticity in SCLC^41^. Predicted Tuft state drivers include *Ascl2, Pou2f3, Gng13, Avil, Rgs13,* and other tuft-specific genes (Fig. 4j), with ASCL2 recently implicated in the POU2F3^+^ tuft-state of prostate neuroendocrine cancer^4^. This analysis also uncovered drivers of various TN^-^ populations (Extended Data Fig. 6e,f and Supplementary Tables 3,5): Wnt-related genes (*Tcf7l2, Lrp1), Notch1, Fgfr1, Vim,* and *Sox9*^105,106^ promoted the Cluster 5, mesenchymal/fibroblast state. Early enrichment of *Elf3, Krt19,* and *Epcam* followed by induction of ionocyte markers *Atp6v0e* and *Cftr* drove fate bias to Cluster 8—a state enriched for epithelial, luminal hillock^107^, and ionocyte markers. *Twist1, Mdk*, and *Fzd2* induction were associated with fate switching towards TN^-^ states enriched for proliferative and embryonic stem markers. Altogether, analyses of CellTagged RPM and RPMA allografted tumour cells highlight remarkable plasticity between SCLC-A, -N, and -P fates at unprecedented single-cell resolution and shed light on both established and potential new drivers of tumour cell fate decisions.

### PTEN loss promotes POU2F3 in basal-derived SCLC

Our data suggest genetic alterations (*Myc* gain, *Ascl1* loss) and cell of origin function together to influence SCLC fate and plasticity. In addition to increased *MYC* and decreased *ASCL1* in human SCLC-P versus other subtypes (Fig. 5a), genomic data demonstrate *PTEN* alterations are enriched in *POU2F3*^high^ tumours (∼42% vs ∼8% in *POU2F3*^low^)^58^. Similarly, analysis of copy number data from n=112 human tumours^38^ shows that *PTEN* deletions occur in ∼63% (10/16) of *POU2F3*^high^ and only 19% (18/96) of *POU2F3*^low^ tumours (Fisher’s exact test, ***p < 0.0006) (Fig. 5a). Thus, we predicted that *PTEN* loss may promote SCLC-P fate and/or provide a selective advantage for SCLC-P growth.

**Fig. 5:**
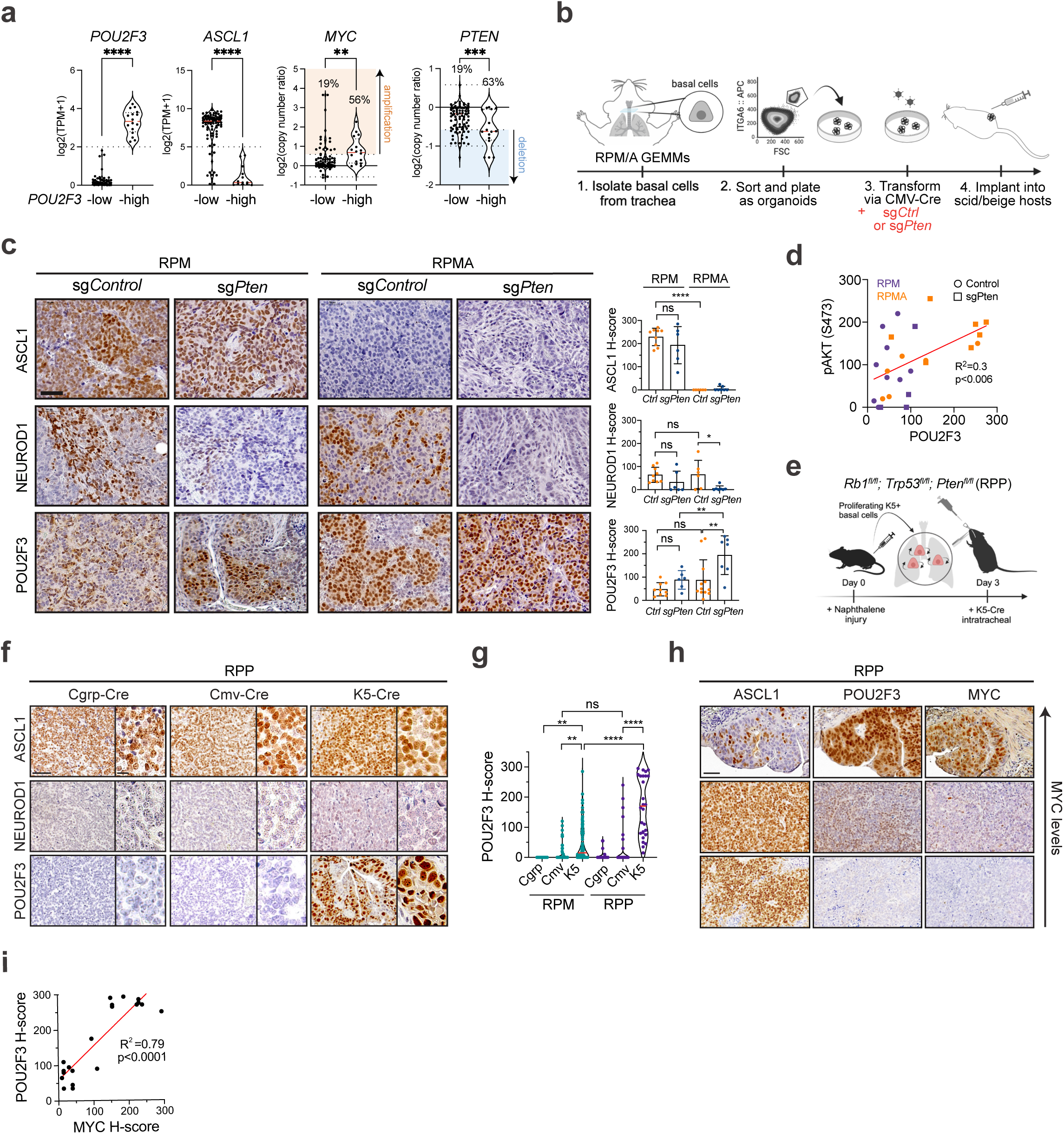
PTEN loss and MYC cooperate to drive POU2F3^+^ basal-derived SCLC. See also Extended Data Figs. 7 and 8. **(a)** mRNA expression of *POU2F3* and *ASCL1* as log2(TPM+1) and copy number data for *MYC* and *PTEN* as log2(copy number ratio) in n=112 human SCLC tumours^38^ grouped by *POU2F3* status (n=96 *POU2F3-*low, n=16 *POU2F3-*high). Percent of tumours with *MYC* amplification (>log2(3/2)=0.58) and *PTEN* heterozygous (<log2(1/2)=-0.58) or deep deletion (< -1) indicated. Median (dashed red line) and upper and lower quartiles (dotted lines) are indicated. Student’s unpaired t-tests on gene expression data and Fisher’s exact tests on copy number data. **** p<0.0001, *** p<0.001, ** p<0.01. **(b)** Schematic depicting generation of RPM and RPMA basal organoids and allografts with loss of *Pten*. **(c)** Representative IHC images from RPM and RPMA basal-organoid-derived parental or LCV2-*sgCtrl* tumours compared to tumours with LCV2-*sgPten* for indicated markers (left). H-score IHC quantification for indicated proteins (right). Each dot represents one tumour. For each genotype, n=3-10 parental, n=2-6 sg*Ctrl,* and n=6-9 sg*Pten* tumours were quantified. Parental and sg*Ctrl* tumours are grouped in the same category labeled as *Ctrl.* **(d)** Simple linear regression analysis of H-score quantification of phospho-AKT (pAKT S473) versus POU2F3 in RPM and RPMA parental and sg*Ctrl* (Control) and sg*Pten* basal allograft tumours, coloured by genotype (RPM=purple, RPMA=orange). Goodness of fit (R^2^) and p-value indicated. **(e)** Schematic depicting method to induce SCLC from basal cells in the *Rb1^fl/fl^;Trp53^fl/f^;Pten^fl/fl^* (RPP) GEMM. **(f)** Representative IHC for SCLC subtype markers on RPP tumours initiated by indicated Ad-Cre viruses. Scale bar=10 μm on high-magnification inset. **(g)** Violin plot of H-score quantification of POU2F3 IHC in RPM versus RPP GEMM tumours initiated by indicated Ad-Cre viruses. Each dot equals one tumour. N=11-132 tumours quantified per Ad-Cre group per genotype from n=4-19 mice per condition. Median (red bar) and upper and lower quartiles (solid line) are indicated. **(h)** Representative IHC images on serial sections of RPP K5-Cre tumours for indicated markers on MYC-high, -medium, and -low regions (Y-axis). **(i)** Simple linear regression analysis of H-score quantification of MYC versus POU2F3 IHC in RPP K5-Cre tumours. N=21 tumours quantified from n=4 mice. Goodness of fit (R^2^) and p-value indicated. All scale bars=50 μm unless otherwise noted. Unless otherwise noted, statistical tests are two-way ANOVA. **** p<0.0001, ** p<0.009, * p<0.02, ns=not significant p>0.05. Error bars represent mean ± SD.

To address the consequence of *Pten* loss in basal-derived SCLC, CRISPR-mediated knockout of *Pten* or a non-targeting sgRNA (sg*Ctrl*) was introduced in RPM and RPMA basal organoids (Fig. 5b). After validating *Pten* editing and downstream induction of phospho-AKT (Extended Data Fig. 7a,b), basal organoids were implanted into scid/beige mice to generate allografts. Consistent with its known tumour suppressive role, *Pten* loss expedited growth of both RPM and RPMA basal allografts (Extended Data Fig. 7c). Analysis of resulting allografts confirmed the expected loss of ASCL1 and gain of YAP1 in RPMA allografts compared to RPM controls, revealing no change in ASCL1 or YAP1 levels upon *Pten* loss (Fig. 5c and Extended Data Fig. 7d). However, *Pten* loss led to a striking increase in POU2F3 in both RPM and RPMA allografts, with some RPMA tumours having robust POU2F3 in nearly 100% of cells (Fig. 5c). POU2F3 levels significantly correlated with phospho-AKT levels (Fig. 5d and Extended Data Fig. 7d). Enrichment of POU2F3 in *sgPten* versus *sgCtrl* tumours coincided with a notable reduction in NEUROD1, particularly in RPMA tumours (Fig. 5c). Thus, PI3K/AKT pathway activity preferentially promotes formation of, or evolution toward, SCLC-P, seemingly at the expense of SCLC-N.

Pathological review confirmed that compared to RPM and RPMA controls, *Pten-*deleted RPMA tumours exhibited increased histopathological heterogeneity—including regions of adeno-, adeno-squamous, and squamous cell carcinoma enriched for KRT5 and DNP63 expression (Extended Data Fig. 7e). Within individual tumours, RPM control, RPM *Pten*-deleted, and RPMA control tumours were predominantly SCLC, while each RPMA *Pten-*deleted tumour comprised ∼40% SCLC and ∼60% non-SCLC regions— most often adeno-squamous (Extended Data Fig. 7f). Consistent with these observations, squamous tumours are known to derive from basal cells, and human squamous tumours exhibit activation of the PI3K/AKT pathway and MYC^79,108,109^, altogether suggesting ASCL1 status is a master determinant of basal cell fate decisions between squamous and neuroendocrine lineages in the presence of MYC and AKT. RPMA-sg*Pten* models that develop both tuft-like and squamous-like tumours suggest a lineage relationship between these histopathologies. The connections between SCLC-P and squamous lung cancer are currently unknown, but a recent study detected human SCLC-P near squamous cell carcinoma in cases of combined SCLC^110^, suggesting a potential transition between these histologies. Taken together, the basal cell exhibits a remarkable capacity to generate the major distinct lung cancer subtypes in accordance with precise genetic alterations, and the RPMA-sg*Pten* model enables further exploration of these lineage relationships and mechanisms.

### PTEN loss and MYC cooperate to drive POU2F3^+^ SCLC

Given *Pten* loss promoted SCLC-P in RPM and RPMA basal-derived allografts, we questioned whether loss of *Pten* in autochthonous GEMMs may also promote the tuft-like state of SCLC from a basal origin. In the *Rb1^fl/fl^/Trp53^fl/fl^/Pten^fl/fl^*(RPP) GEMM^111,112^, tumours initiated from NE cells lack MYC and resemble *Mycl*-high SCLC-A, without expression of NEUROD1 or POU2F3^68,111,113^. To test if *Pten* loss promotes SCLC-P from basal cells in the RPP model, tumours were initiated with K5-Cre following naphthalene injury and compared to CGRP-Cre and CMV-Cre-initiated tumours (Fig. 5e–g). K5-initiated tumours developed in RPP GEMMs with a latency of 145 days (Extended Data Fig. 8a), comparable to the latency observed when tumours are initiated from NE cells in this model (n=164 days)^68^. Similar to findings from NE cells of origin, basal-derived RPP tumours highly expressed ASCL1 and lacked NEUROD1 (Fig. 5f and Extended Data Fig. 8b). Yet, in contrast to NE cells of origin, K5-derived RPP tumours highly expressed POU2F3—significantly more than both CGRP- and CMV-Cre initiated tumours from the same model (Fig. 5g). Individual POU2F3^+^ tumours in the RPP GEMM often harboured ASCL1^+^ tumour cells (Fig. 5h), suggesting a basal origin permits neuroendocrine–tuft plasticity in RPP SCLC. Thus, *Pten* loss promotes SCLC-P in basal cells in both allograft and autochthonous models under multiple genetic conditions.

PTEN loss and PI3K/AKT pathway activation can upregulate MYC via multiple mechanisms in cancer^114,115^. Consistent with previous reports^58^, analysis of the largest publicly available SCLC dataset with matching transcriptomic and copy number data reveals that *MYC* is amplified in ∼56% (9/16) of *POU2F3*^high^ and only 19% (18/96) of *POU2F3*^low^ tumours (Fisher’s exact test, **p < 0.003) (Fig. 5a). In this dataset, co-occurring *PTEN* and *MYC* copy number variation were detected in 25% (4/16) of *POU2F3*^high^ tumours, and only 7.3% (7/96) of *POU2F3*^low^ tumours (Fisher’s exact test, p = 0.05). Given *MYC*’s selective overexpression and amplification in SCLC-P and co-occurrence with *PTEN* loss in human tumours, we assessed MYC levels in basal-derived RPP tumours. Indeed, MYC was detected in 100% of POU2F3^+^ RPP tumours, and its expression level highly correlated with POU2F3 intensity (Figs. 5h,i). In contrast, MYC was absent in uniformly ASCL1^+^ RPP tumours (Fig. 5h), suggesting a tuft-lineage-specific tolerance for MYC in the RPP model. Notably, POU2F3 was significantly enriched in RPP tumours from a basal origin, even when compared to MYC-driven RPM tumours from a basal origin (Fig. 5g). Together, these data strongly suggest that activation of PI3K/AKT signaling acts upstream of MYC to cooperatively facilitate tuft fate in SCLC, more so than high MYC alone. Moreover, AKT activation through either *PTEN* deletion or expression of exogenous myristoylated-AKT leads to upregulation of MYC in human POU2F3^+^ SCLC cells (Extended Data Fig. 8c,d). We speculate that compared to neuroendocrine cells, SCLC arising from basal cells have increased tolerance, and/or transcriptional flexibility, in response to oncogenic MYC signaling, permitting tuft lineage conversion. A key role for PI3K/AKT signaling in promoting SCLC-P (and potentially suppressing SCLC-N) represents a previously unappreciated driver that could potentially be therapeutically exploited.

### Human SCLC harbours an inflammatory basal-like subset

We next sought to determine whether a basal-like state exists in human SCLC, and to determine the concordance between basal-derived murine SCLC and human tumours. SCLC lacking expression of A, N, and P have been previously classified as YAP1^+^, inflamed, or mesenchymal^12,13,36,38–40^, but these “lineage-negative” tumours remain controversial and how they relate to other SCLC subtypes has been unclear.

Whether the “lineage-negative” tumours are enriched for signatures of basal cells and their derivatives is similarly unknown. To address these issues, we analyzed bulk transcriptomic data from the largest real-world SCLC dataset to date comprising 944 human biopsies.

Based on the relative expression of lineage-defining transcription factors (evaluated as Z-scores), human samples were categorized as A, N, or P if they exhibited high expression (positive Z-score) exclusive to one gene, ‘Mixed’ if high expression was observed for any combination of A/N/P genes, or ‘Lineage-negative’ (Lin^-^) if they exhibited low expression of all three (Fig. 6a). Application of a normal Basal cell signature^14^ including basal-specific genes *TP63, KRT5, ICAM1,* and *EPAS1* revealed enrichment particularly in Lin^-^ tumours (Fig. 6a, Extended Data Fig. 9a, and Supplementary Table 2). Interestingly, the Basal cell signature and basal genes were also enriched in a small subset of each of the other tumour classifications (i.e, A, N, P, Mixed) (Fig. 6a). The distribution of basal phenotypes across human SCLC subsets is consistent with basal-like tumour cells representing a minor tumour subpopulation across many samples (as we observe in mouse model data).

**Fig 6:**
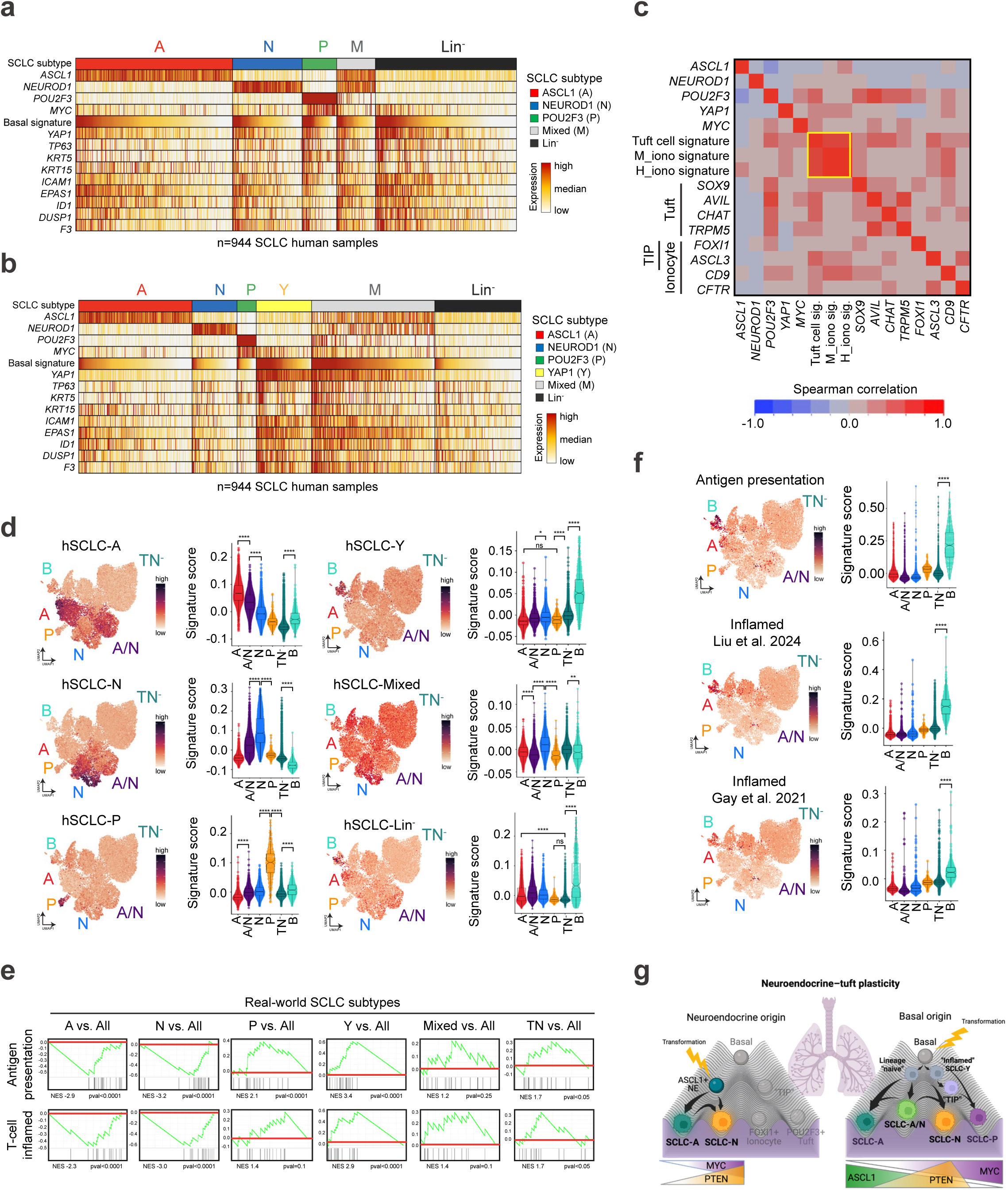
Human SCLC harbours basal-like subset. See also Extended Data Fig. 9 and Supplementary Tables 2 and 6. **(a)** Heatmap displays expression by bulk RNA-seq of lineage-related transcription factors, a basal cell signature, and basal markers in n=944 human SCLC biopsies, grouped and annotated by subtype. **(b)** Heatmap as in Fig. 6a with expression by bulk RNA-seq of lineage-related transcription factors, a basal cell signature, and basal markers, grouped and annotated by subtype including a YAP1 subtype. **(c)** Spearman correlation matrix depicting individual gene or gene signature correlations by bulk RNA-seq in n=944 human SCLC biopsies (Supplementary Table 2). Data include subtype markers, *MYC*, and annotated Tuft, tuft-ionocyte-progenitor (“TIP”), and ionocyte markers. Yellow box indicates tuft and ionocyte signatures with the strongest correlations. **(d)** Expression of human subtype signatures (hSCLC-A, -N, -P, -Y, -Mixed, -TF^-^) derived from the real-world bulk RNA-seq dataset applied to RPM and RPMA basal allograft tumour cells from Fig. 3e in UMAP space (left) or per assigned SCLC fate in the form of a violin plot (right) (Supplementary Table 6). **(e)** GSEA for established Antigen presentation and T-cell inflamed signatures^12^ in human real-world subtypes as indicated. **(f)** Antigen presentation or “Inflamed” SCLC tumour signatures derived from bulk RNA-seq and/or proteomics data on human SCLC tumours and applied to RPM and RPMA basal allograft tumour cells from Fig. 3e in UMAP space (left) or per assigned SCLC fate in the form of a violin plot (right) (Supplementary Table 6). **(g)** Graphical abstract depicting SCLC fate trajectories possible from a neuroendocrine (left) vs basal (right) cell of origin. Thickness of arrows indicates frequency that trajectories are likely to occur in the RPM GEMM. MYC, PTEN, and ASCL1 levels vary with fate according to annotations below the Waddington landscapes. Box-whisker overlays on all violin plots indicate median and upper and lower quartile. One-way ANOVA with Tukey’s correction. **** p<0.0001, ** p<0.0004, * p<0.004, ns=not significant, p>0.05.

The Basal signature was highly correlated with *YAP1* expression in human tumours—particularly in the Lin^-^ tumours, but also across subtype classifications (Fig. 6a and Extended Data Fig. 9a). Due to the correlation of *YAP1* with basal signatures, tumours were next classified into six groups—this time including YAP1 as a subtype classification if samples exhibited exclusively high *YAP1* (positive Z-score) (Fig. 6b). Following this categorization, enrichment of the Basal signature extended beyond the YAP1-classified tumours, with particularly strong expression in Mixed tumours, and a relative depletion in the remaining Lin^-^ tumours (Fig. 6b). Further study is necessary to characterize the phenotype of the non-basal/Lin^-^ tumours. We speculate that enrichment of the Basal signature in Mixed tumours may be an indication that the basal state facilitates subtype diversity, consistent with our findings of an A/N hybrid state enriched in basal-derived murine tumours (Fig. 1i,j and 4c). The presence of a basal-like identity in YAP1 and Mixed tumours argues that the majority of the previously described YAP1^+^ subset of human SCLC are most likely enriched for basal-like tumour cells.

In addition to neuroendocrine and tuft cells, normal lung basal cells can spawn rare CFTR^+^ ionocytes and recently discovered tuft-ionocyte progenitor (“TIP”) cells marked by co-expression of tuft- and ionocyte-lineage markers, including *POU2F3* and *FOXI1* or *ASCL3,* respectively^14,18,71,116^. The ionocyte marker, *FOXI1,* has been detected in other tuft-like tumours across tissues^11,38,65^. Consistently, we identified a strong correlation between normal tuft and ionocyte genes and signatures^14^ within human SCLC tumours (Fig. 6c, Extended Data Fig. 9b, and Supplementary Table 2). When comparing each human subtype to all other subtypes, gene set enrichment analyses (GSEA) demonstrate human SCLC-A is enriched for neuroendocrine cell signatures, SCLC-Y is enriched for basal cell signatures, and SCLC-P is enriched for both tuft and ionocyte cell signatures (Extended Data Fig. 9c). While ionocyte signatures were most enriched in SCLC-P, some enrichment also occurred in SCLC-A (Extended Data Fig. 9c), consistent with observations of overlapping marker expression between normal ionocytes and neuroendocrine cells^14,77^. Human subtype signatures derived from the real-world dataset remarkably parallel the fates observed in murine basal-derived SCLC tumours (Fig. 6d), including enrichment for the SCLC-Y signature in the Basal-like state (Fig. 6d), and enrichment of Ionocyte signatures in A, P, and basal-like states (Extended Data Fig. 9d). Together, these data further demonstrate that the basal cell can generate tumours that recapitulate the spectrum of SCLC heterogeneity, and that SCLC subtypes mirror the basal cell and its rare basal-derived lineages^14^. Going forward, it will be important to determine whether tuft and/or TIP cells can give rise to SCLC, and if so, the extent of SCLC plasticity from these cells of origin.

In human SCLC, YAP1^+^ tumours are observed to be more prevalent in limited-versus extensive-stage disease, suggesting they may arise earlier, and have a better prognosis^39^—perhaps due to increased inflammation^12,13,36,38–40^ and anti-tumour immune responses. Indeed, compared to other subtypes, the real-world SCLC-Y subset that is enriched for basal-marker expression is significantly enriched for established “T-cell Inflamed” and “Antigen Presentation” signatures^12^ (Fig. 6e). Consistently, in the mouse basal tumour state, the Antigen Presentation signature and two additional human “Inflamed” SCLC signatures^12,38^ are significantly enriched (Fig. 6f). These data suggest the basal-like state of SCLC has an inflammatory nature, and that high basal-signatures in human SCLC may function as a prognostic indicator of immunotherapy response. Moreover, signals that promote the basal state may have the capacity to warm up the “immune-cold” SCLC landscape.

The strong concordance of human subtype signatures within our basal-derived SCLC mouse models (Fig. 6d) underscore the basal cell of origin and basal cell differentiation as drivers of SCLC development and plasticity. Specifically, our findings suggest a model whereby the basal cell gives rise to all established SCLC subtype states, including SCLC-A, -N, -P, “Mixed”, and “TIP”-like phenotypes (Fig. 6g). Coupled with the abundance, location, and capacity of normal basal cells to proliferate and differentiate into the major SCLC lineages, our findings strongly suggest the basal cell as an unappreciated but highly likely cell-of-origin for SCLC and potentially for other tumours exhibiting neuroendocrine–tuft heterogeneity.

It will be important to determine if other tumours that exhibit neuroendocrine–tuft heterogeneity arise from a basal origin and/or co-opt a basal-like state to achieve lineage plasticity. Recently, we established that the globose basal cell of the olfactory epithelium is an origin for olfactory neuroblastomas that have both neuroendocrine and tuft/microvillar states^6^. Additionally, recent work has implicated a basal intermediate state in the conversion of lung^117^ and prostate adenocarcinoma^4,26^ to neuroendocrine tumours during resistance to targeted therapy. Thus, SCLC can likely arise *de novo* in basal cells, and/or dedifferentiate to a basal-like state during therapeutic stress. Interestingly, in models of adenocarcinoma-to-SCLC transdifferentiation, MYC permits cells to pass through a critical bottleneck to achieve lineage plasticity^117^. In a recently developed model of pancreatic cancer progression, MYC is similarly required to drive tuft-to-neuroendocrine lineage conversion^118^. Coupled with our findings here, this suggests the basal state may cooperate with MYC signaling to govern lineage plasticity across multiple cancer types and contexts.

Altogether, we establish the basal cell as a plausible origin for the highly plastic nature of SCLC. We develop new POU2F3-enriched lung tumour models—including immune-competent models—that are amenable to genetic, microenvironmental, and therapeutic studies. Recent studies suggest the mammalian SWI/SNF complex as a therapeutic dependency in SCLC-P^31,41^. Further work is necessary to ascertain mechanisms leading to increased PI3K/AKT signaling in SCLC-P and whether it represents a therapeutic vulnerability. These immune-competent GEMMs will be important resources for drug testing and for understanding the inflammatory basal state. Likewise, the organoid-derived SCLC allograft models characterized here are remarkably similar to autochthonous tumours but have additional flexibility for genetic manipulation: facilitating lineage-tracing, genetic screens, and easier pharmacological studies than traditional GEMMs. The new organoid-based models will be valuable to better understand the biology governing neuroendocrine–tuft plasticity and test the utility of plasticity-directed therapies for cancer treatment. We expect that discoveries using these models will shed light on plasticity mechanisms that are relevant to SCLC and many other tumours with neuroendocrine–tuft-like states^4–8,24–27,118,119^.

## Methods

### Resource availability

#### Materials availability

There are limitations to the availability of basal-derived organoid lines generated in this study due to their derivation from primary cells in the tracheal epithelium. Human SCLC tissue used in this study is not available due to sample scarcity. Human transcriptomics data from Caris Life Sciences used for this study are not publicly available but can made available upon reasonable request. The deidentified sequencing data are owned by Caris Life Sciences, and qualified researchers can apply for access by signing a data usage agreement.

#### Data and code availability

All single-cell RNA-seq data have been deposited at GEO (GSE279200) and will be publicly available on the date of publication. All original code has been deposited on GitHub (https://github.com/TOliver-Lab/Ireland_Basal_SCLC_2024) and available on the date of publication. Any additional information required to reanalyze the data reported in this paper is available from the lead contact upon request.

### Experimental models and study participant details

#### Mice

*Rb1^fl/fl^;Trp53^fl/fl^;H11b-LSL-Myc^T58A/T58A^-Ires-Luciferase* (RPM) (JAX #029971)^68^, *RPM;Ascl1 ^fl/fl^* (RPMA)^43^, *Rb1^fl/fl^;Trp53^fl/fl^;Rbl2/p130^fl/fl^*(RPR2)^85^, and *Rb1^fl/fl^;Trp53^fl/fl^;Pten^fl/fl^*(RPP)^111,112^ mice have been previously described. SCID/beige mice (CBSCGB) are purchased and available from Taconic and Charles River Laboratories.

All mice were housed and treated according to regulations set by the Institutional Animal Care and Use Committee of Duke University. Viral infections were performed in a Biosafety Level 2+ room following guidelines from Duke University Institutional Biosafety Committee. Male and female mice were distributed equally for all experiments.

#### Basal-derived organoid cultures and cell lines

Basal-derived organoid cultures from RPM, RPMA, and RPR2 mice were obtained and transformed ex vivo (see Method details). Organoid lines were determined to be free of pathogens by IDEXX 18-panel mouse pathogen testing and confirmed mycoplasma-negative before implantation to SCID/beige hosts. Cell lines used in this study include HEK 293T/17 cells (ATCC cat# CRL-11268) to produce lentivirus and H1048 SCLC cells (ATCC cat# CRL-5853). Cell lines were tested for mycoplasma every three months and were negative. Cell line identities were confirmed via STR profiling within six months of usage, last performed in July of 2024.

#### Patient tissue for immunostaining

Biopsies for establishment of patient-derived xenograft (PDX) models were performed following written informed consent from patients under an Institutional Review Board-approved protocol at Memorial Sloan Kettering (IRB14-091). Models were established and characterized as previously described^120^. As previously described^42^ for human biopsies from Huntsman Cancer Institute (HCI), all patients provided informed consent for the collection of specimens, approved by the University of Utah Institutional Review Board (IRB_00010924 and IRB_00089989) in accordance with the U.S. Common Rule. For tissue micro-arrays, human biopsies collected at Washington University St. Louis were acquired with approval under IRB_202008098.

#### Caris Life Sciences patient cohort

Caris real-world data derive from a retrospective review of patient tumour specimens (n=944) with a diagnosis of small cell lung cancer (based on pathological confirmation by local pathologists) submitted to a Clinical Laboratory Improvement Amendments (CLIA)-certified laboratory (Caris Life Sciences) for molecular profiling. The present study was conducted in accordance with the guidelines of the Declaration of Helsinki, Belmont Report, and US Common Rule and in compliance with policy 45 CFR 46.101(b). The study was conducted using retrospective, de-identified clinical data, and patient consent was therefore not required.

### Method details

#### Naphthalene injury model and tumour initiation in mice

Mice at 6–8 weeks of age were treated intraperitoneally (i.p.) with 275 mg/kg naphthalene before 9AM in corn oil as described^121^, 72 h before administration of adenoviral Cre—a timepoint where KRT5^+^ basal cells are shown to be abundant and proliferative^122^. Following naphthalene treatment, mice were infected by intratracheal (RPM and RPP) or intranasal (RPP) instillation with 1×10^8^ plaque-forming units of Ad5-K5-Cre adenovirus (University of Iowa cat# VVC-Berns-1547) using established methods^123^. No observed differences in latency or tumour phenotype occurred in RPP mice with intratracheal versus intranasal inoculation methods, so both were included in results. Briefly, mice were anesthetized with isoflurane at a flow rate of 20–25 ml/h, depending on the size and sex of the mouse. The optimal breathing rate was approximately one breath every 2–3 sec. For intratracheal instillation, mice were positioned on a platform with their chest hanging vertically beneath them. A steel feeding tube or Exel Safelet IV catheter (needle removed) was slid into the trachea, and 63 μl of viral cocktail consisting of 10 mM CaCl_2_ (Sigma cat# C5670) +1×10^8^ pfu adenovirus + MEM (Thermo cat#11095080) up to 63 μL was administered via a P200 pipette to the catheter opening. Mice were maintained in this position until the entire volume was dispensed, then were monitored until they regained full motility and recovered from anesthesia. For intranasal instillation, mice were held in a supine position and administered 63 μL of identical viral cocktail via a P20 pipette, alternating between the left and right naris for each drop.

Administration of other Ad-Cre viruses (CGRP-, cat# VVC-U of Iowa-1160; SPC-, cat# VVC-U of Iowa-1168; CCSP-, cat# VVC-U of Iowa-1166; CMV-, cat# VVC-U of Iowa-5) also occurred in mice 6–8 weeks of age with identical methods, intratracheally, but without naphthalene injury, as previously described^43,68,113^.

#### MicroCT imaging

To monitor tumour development in autochthonous models, mice were imaged beginning four weeks after Ad-Cre administration for RPM mice, 8 weeks for RPP mice, and every two weeks thereafter. Mice were anesthetized with isoflurane and imaged using a small animal Quantum GX2 microCT (Perkin Elmer).

Quantum GX2 images were acquired with 18 sec scans at 45 μm resolution, 90 kV, with 88 mA of current. Mice were sacrificed when tumour burden resulted in any difficulty breathing or significant weight loss.

#### Immunohistochemistry (IHC)

For IHC of autochthonous mouse models, lungs were inflated with 1X PBS, extracted, and individual lung lobes and trachea were collected for fixation. Tissues were fixed in 10% neutral buffered formalin for 24 h at room temperature (RT), washed in PBS and transferred to 70% EtOH. Formalin-fixed paraffin embedded (FFPE) sections at 4–5 μm were dewaxed, rehydrated, and subjected to high-temperature antigen retrieval by boiling 20 min in a pressure cooker in 0.01 M citrate buffer at pH 6.0. Slides were quenched of endogenous peroxide in 3% H_2_O_2_ for 15 min, blocked in 5% goat serum in PBS/0.1% Tween-20 (PBS-T) for 1 h, then stained overnight with primary antibodies in blocking buffer (5% goat serum or SignalStain antibody diluent, Cell Signaling Technology, cat#8112). For non-CST primary antibodies, an HRP-conjugated secondary antibody (Vector Laboratories) was used at 1:200 dilution in PBS-T, incubated for 45 min at RT, and followed by DAB staining (Vector Laboratories). Alternatively, CST primary antibodies were detected using 150 μL of SignalStain Boost IHC Detection Reagent (CST cat#8114). All staining was performed with Sequenza cover plate technology. The primary antibodies include: ASCL1 (Abcam cat#211327) 1:300; NEUROD1 (Abcam cat#109224) 1:300; POU2F3 (Sigma cat# HPA019652) 1:300; YAP1 (CST cat#14074) 1:300; HES1 (CST cat#11988) 1:300; DNP63 (R&D cat# AF1916) 1:400; phospho-AKT Ser473 (CST cat#4060) 1:100; NKX2-1 (Abcam cat# ab76013) 1:250; and KRT5 (BioLegend cat#905501) 1:1000. For manual H-score quantification, whole slides were scanned in using a Pannoramic Midi II automatic digital slide scanner (3DHistech) and images were acquired using SlideViewer software (3DHistech). IHC quantification from primary tumour models included tumours from both the trachea and lung lobes. H-score was quantified on stained slides on a scale of 0–300 taking into consideration percent positive cells and staining intensity as described^124^, where H score = % of positive cells multiplied by intensity score of 0–3. For example, a tumour with 80% positive cells with high intensity of 3 has a 240 H-score.

#### Immunofluorescence

Lung and tumour tissue was collected and fixed for at least 24 h in 10% neutral buffered formalin, then transferred to 70% EtOH before embedding in paraffin. Wildtype and transformed organoids (>1×10^6^ cells) were collected in ∼0.5–1 mL of organoid media with a P1000 pipette tip, then transferred to a conical tube with 10 mL of 10% formalin. Organoids were fixed at room temperature in formalin for 24 hrs. After fixation, organoids were spun down at 500xg for 5 min, then washed in 70% EtOH. EtOH was removed and organoids were resuspended in ∼300 μL of 3% low melting agarose gel (microwaved to melt, then incubated in a 50°C water bath for 30 min) with a wide-bore P1000 pipette tip, then transferred to one well of a 96-well V-bottom plate. When agarose solidified (∼3–5 min, RT), agarose plugs containing organoids were transferred from the well-plate to histology cassettes, placed in 70% EtOH, then subject to FFPE and sectioning for slides. Prior to staining, slides were rehydrated in CitriSolv 2x 3 min, 100% EtOH 2x 3 min, 90% EtOH 1x 3 min, 70% EtOH 1x 3 min, 40% EtOH 1x 3 min, and dH20 1x 5 min. Rehydrated tissue was subject to high-temperature antigen retrieval by boiling 15 min in a pressure cooker in 0.01 M citrate buffer at pH 6.0. Slides were cooled at room temperature for 2 h and positioned for staining in Sequenza staining racks (Thermo Scientific cat#10129-584). Slides were blocked at RT for 1 h in 10% donkey serum in PBS/0.2% Tween-20 (PBS-T). For primary mouse-on-mouse tissue (M.O.M.) staining, M.O.M. IgG Blocking Reagent (Vector Labs cat# PK-2200) was also added according to manufacturer’s protocol. Primary antibody was diluted in 10% donkey serum in PBS-T, added to slides, and slides were incubated overnight at 4°C. The following day, slides were washed 3x with PBS-T, then stained with secondary antibody diluted in 10% donkey serum. For M.O.M. staining, M.O.M. Protein Concentrate (Vector Labs cat# PK-2200) was added to the secondary antibody solution, according to manufacturer’s protocol. Slides were then subject to 3x additional washes with PBS-T, followed by DAPI staining (1 μg/mL in PBS-T) for 20 min. Following 3x additional washes in PBS-T, slides were coverslipped with Aqua-Poly/Mount mountant (Polysciences Inc cat#18606-20). Primary antibodies included: anti-mouse ASCL1 (BD Pharmingen cat#556604) 1:25; anti-rabbit ASCL1 (Abcam cat#211327) 1:100; anti-goat NEUROD1 (R&D Systems cat# AF2746) 1:50; anti-rabbit NEUROD1 (Abcam cat#109224) 1:200; POU2F3 (Sigma cat# HPA019652) 1:100; anti-rabbit CCSP/SCGB1A1 (Millipore Sigma cat#07-623) 1:75; anti-rat KRT8 (DSHB cat# TROMA-I) 1:100; anti-mouse FOXJ1 (eBioscience cat#14-9965-80) 1:100; anti-mouse KI67 (BD Pharmingen cat# BDB556003) 1:100; anti-goat DNP63 (R&D cat# AF1916) 1:40. Secondary antibodies were all used at a concentration of 10 μg/mL and included: Donkey anti-rabbit AF488 (Invitrogen cat# A21206); Donkey anti-rat AF568 (Invitrogen cat# A78946); Donkey anti-rat AF647 (Invitrogen cat# A78947); Donkey anti-mouse AF647 (Invitrogen cat# A31571); Donkey anti-goat AF594 (Invitrogen cat# A11058). Slides were imaged on an EVOS M5000 (Invitrogen cat# AMF5000) digital inverted benchtop microscope.

#### Human SCLC cell infections

Human SCLC cell line H1048 was obtained from ATCC and cultured in RPMI media supplemented with 10% fetal bovine serum (FBS), 1% L-glutamine, and 1% penicillin/streptomycin antibiotic cocktail. To generate H1048 sg*NTC,* sg*PTEN,* and myrAKT cell lines, cells were infected with a non-targeting sgRNA (sg*NTC*) or an sgRNA against *PTEN* (sg*PTEN*: *5’-*GAC TGG GAA TAG TTA CTC CC -3’) in the LCV2-hygro backbone (Addgene Plasmid #98291), or infected with the pHRIG-AKT1 lentiviral construct (Addgene, cat# 53583). In brief, high-titer virus (∼1-5 ×10^7^ pfu) was produced using HEK-293T cells transfected with a three-plasmid system including the targeting construct and lentiviral packaging plasmids pCMV delta R8.2 (Addgene Plasmid #8455) and pCMV-VSVG (Addgene Plasmid #8454).

Viruses were harvested at 48 and 72 h post-transfection, concentrated by ultracentrifugation (25,000 RPM for 1.45 h), resuspended in 1X sterile PBS, and stored at -80°C until use. H1048 cells were subject to spinoculation at 37°C, 900 x g, for 30 min. During spinoculation, 0.5–1 million cells per well of a 6-well plate were cultured with 2 mL RPMI, 25 μL HEPES buffer (Thermo Fisher cat# 15630080), 8 μg/mL polybrene (Santa Cruz cat# sc-134220), and 25 μL high-titer virus. Cells were selected 48 h after spinoculation with hygromycin (sgCRISPR) or sorted for GFP to enrich for cells infected with pHRIG-AKT1.

#### Immunoblotting

For human cell line and mouse tumour western blots, protein lysates were prepared as described^68,125^, separated via SDS-PAGE, and transferred with to PVDF membranes (BioRad cat# 1704157) using a Trans-Blot Turbo Transfer System (BioRad cat#1704150). Membranes were blocked for 1 h in 5% milk followed by overnight incubation with primary antibodies at 4°C. Membranes were washed for 3x 10 min at RT in TBS-T. Mouse and rabbit HRP-conjugated secondary antibodies (Jackson ImmunoResearch, 1:10,000) were incubated for 1 h in 5% milk at RT followed by washing 3x 10 min at RT in TBS-T. Membranes were exposed to WesternBright HRP Quantum substrate (Advansta cat# K-12045-D50) and detected on Hyblot CL film (Denville Scientific Inc). Primary antibodies included: ASCL1 (1:1000; Abcam cat#211327); NEUROD1 (1:1000; CST cat#62953); MYC (1:1000; CST cat#5605); PTEN (1:1000; CST cat#9559); POU2F3 (1:1000; Sigma cat# HPA019652), pAKT (Ser473) (1:1000; CST cat#4060), pAKT (Thr308) (1:1000; CST cat#13038), total AKT (1:1000, CST cat#9272), and HSP90 (1:1000, CST cat#4877) as loading control.

#### Basal organoids

##### Tracheal basal cell isolation and organoid culture

Live, normal tracheal basal cells were isolated from RPM, RPR2, and RPMA mice (not treated with Ad-Cre) and grown as organoids as described previously^16,126^. In brief, mice were euthanized and 3-4 tracheas per genotype were isolated in cold DMEM/F12-Advanced media (Thermo Sci cat#12-634-238 +10% FBS,1% L-glutamine,1% Pen/Strep). Tracheas were opened to expose the lumen using a razor blade and forceps. Each trachea was placed in a 1.5 mL Eppendorf tube in 500 μL Dispase (50U/mL, Corning cat #354235) diluted in HBSS-free media (Thermo Fisher cat#14175-095) to 16U/mL, and incubated at RT for 30 min. After incubation, tracheas were transferred to new Eppendorf tubes containing 500 μL of 0.5 mg/mL DNAse (Fisher Sci cat# NC9709009) diluted in HBSS-free media, and incubated for an additional 40 min at RT. Tracheas from each genotype were pooled in a 10 cm dish containing DMEM/F12-Advanced media and forceps were used to gently pull apart the epithelial layers/sheets from the cartilage of each trachea. Media containing all tracheal epithelial sheets per genotype were transferred to a 15 mL conical tube and centrifuged at 4C, 2000 RPM, for 5 min. Supernatant was removed and the remaining cell pellet was resuspended in 1 mL of TrypLE Express (Invitrogen cat#12604013), then incubated at 37C for 5 min. TrypLE was quenched via addition of 10 mL DMEM/F12-Advanced media, then transferred through a 100 um cell strainer into a 50 mL conical tube. Excess tissue was pushed through the cell strainer gently using a plunger from a syringe. Filtered cells were spun down at 2000 rpm for 5 min at 4C and supernatant was removed. The remaining cell pellet was resuspended in 1 mL of FACS buffer (20 mL PBS, 400 μL FBS, 80 μL 0.5M EDTA), spun down, and then stained in 100 μL of FACS buffer containing 1 μg anti-rat ITGA6/CD49 (eBioscience cat#14-0495-85) primary antibody for 30–60 min on ice. Samples were washed 3x in FACS buffer than resuspended in 100 μL of FACS buffer with 1 μg of secondary antibody Goat anti-rat APC (Biolegend cat#405407) and incubated for 30 min in the dark on ice. Samples were washed 3x with FACS buffer, then stained for 15 min with 1 μg/mL DAPI. After 3x additional washes, cells were subject to FACS and DAPI-/ITGA6+ cells were isolated. Resulting basal cells were resuspended in 100% Matrigel (Corning/Fisher cat# CB-40234C or homemade)—20,000 to 100,000 cells per one 50 μL Matrigel dome, then 50 μL of Matrigel was plated per well of a pre-warmed 24-well plate. After Matrigel solidified at 37C, 500 μL of organoid culture media (OCM) consisting of 50% L-WRN conditioned media^127^ (Sigma cat# SCM105 or homemade), 50% DMEM/F12-Advanced media (supplemented with 10% FBS, 1% L-glutamine, 1% Pen-Strep, 10 ng/mL EGF, Thermo Fisher cat# PHG0311, and 10 ng/mL FGF, Thermo Fisher cat# PHG0369), and 10 μM Y-27632 Rho kinase/Rock inhibitor (MedChem Express cat# HY-10071) was added per well. OCM was changed every 2-3 days and organoids were split and expanded using TrypLE upon confluence.

##### Lentiviral transduction of organoids with CellTag Library V1

The CellTag V1 plasmid library was purchased from Addgene (Plasmid#124591) and amplified according to the published protocol for this technology^96^. Briefly, the plasmid library was transformed using Stellar competent cells at an efficiency of ∼220 cfus per unique CellTag in the V1 library. The library was isolated from E. coli culture via the Plasmid Plus Mega Kit (Qiagen, cat#12981), and assessed for complexity via high-throughput DNA sequencing with the Illumina MiSeq (75-cycle paired-end sequencing v3). Generation of the CellTag Whitelist from sequencing data resulted in 13,836 unique CellTags in the 90^th^ percentile of detection frequency. High titer lentivirus (1.5E7 transducing units (TU)/mL) was generated from the CellTag V1 library following published protocols^42,128^, and titered based on GFP fluorescence with 293T cells (ATCC, cat# CRL11-268).

Normal (no Cre-mediated recombination) RPM, RPMA, and RPR2 basal organoids were expanded for ∼3.5 weeks post-isolation, then ∼1 ×10^6^ cells were transduced with the CellTag V1 lentiviral library. Organoids were dissociated into single cells with TrypLE (Invitrogen cat#12604013) for 30 min and subject to mechanical dissociation every 10 min of TrypLE incubation. TrypLE was quenched and cells were pelleted and resuspended in 500 μL of OCM plus 8 μg/mL polybrene (Santa Cruz, cat# sc-134220) and 25 μL of CellTag V1 high-titer lentivirus, then plated in one well of a 24-well plate. Cells were spinoculated at 300 x g for 30 min at RT to increase transduction efficiency, incubated immediately following spinoculation for 3–6 h at 37C, then pelleted and replated in 50 μL of Matrigel and 500 μL of fresh viral supernatant. Organoids were incubated for 24 h, then viral media was replaced with normal OCM for organoid expansion. GFP was visible in >50% of cells as soon as 24 h after viral transduction.

##### Basal organoid Cre administration

Normal basal organoids, or CellTagged basal organoids expanded for ∼8–10 weeks to allow clonal expansion, were subject to Cre-mediated transformation. Because TAT-Cre treatment resulted in unreliable levels of recombination (Extended Data Figs. 3a, 4g, and 5a), we utilized high titer adenoviral CMV-Cre (U of Iowa, cat# VVC-U of Iowa-5) to recombine all genotypes, including those for CellTagging experiments. Of note, n=4 non-CellTagged RPM basal allograft replicates, successfully recombined with TAT-Cre, were subject to single-cell RNA-seq and included in scRNA-seq data analysis alongside the CellTagged, CMV-Cre RPM recombined allograft. Importantly, no transcriptional or marker expression distinctions between CMV-and TAT-Cre recombined RPM basal organoids or allograft tumours was noted. For all other samples, successful recombination with Ad-CMV-Cre occurred by: 1) dissociating organoids into single cells (∼500k-1M cells) using TrypLE (Invitrogen cat#12604013) for 30 min at 37C with mechanical dissociation every 10 min; 2) spinoculating (300xg, RT, 30 min) organoids in 2.5-5E7 pfu CMV-Cre in 500 μL OCM + 10 μg/mL polybrene in a 24-well plate; 3) incubating 4–6 hours at 37°C; then 4) seeding in Matrigel and propagating as normal organoid cultures, described above. Full recombination of all alleles for each genotype was confirmed by recombination PCR, four weeks post Cre treatment.

##### PCR validation of recombination efficiency

Qiagen DNeasy kit (cat#69506) was used to isolate genomic DNA from basal-derived organoids following exposure to Cre. Fully recombined tumour-derived cell lines from each genotype were used for positive recombination controls. DNA concentrations were measured on a BioTek Synergy HT plate reader. Equal quantities of tumour genomic DNA (100 ng) were amplified by PCR with GoTaq (Promega cat# M7123) using primers to detect *Rb1* recombination: D15’-GCA GGA GGC AAA AAT CCA CAT AAC-3’,1lox5’ 5’-CTC TAG ATC CTC TCA TTC TTC CC-3’, and 3’ lox 5’-CCT TGA CCA TAG CCC AGC AC-3’. PCR conditions were 94 deg 3 min, 30 cycles of (94°C 30 s, 55°C 1 min, 72°C 1.5 min), 72°C 5 min, hold at 4°C. Expected band sizes were ∼500 bp for the recombined *Rb1* allele, and 310 bp for the unrecombined/floxed allele. Primers to detect *Trp53* recombination include the following: A 5’-CAC AAA AAC AGG TTA AAC CCA G-3’, B 5’-AGC ACA TAG GAG GCA GAG AC-3’, and D 5’-GAA GAC AGA AAA GGG GAG GG-3’. PCR conditions were 94°C 2 min, 30 cycles of (94°C 30s, 58°C 30s, 72°C 50s), 72°C 5 min, hold at 4°C. Expected band sizes were 612 bp for the *Trp53* recombined allele, and 370 bp for the unrecombined/floxed allele. Primers to detect *Myc^T58A^*recombination include the following: CAG-F2 5’-CTG GTT ATT GTG CTG TCT CAT CAT-3’, MycT-R 5’-GCA GCT CGA ATT TCT TCC AGA-3’. PCR conditions used were 94 deg 2 min, 35 cycles of (95°C 30 sec, 60°C 30 sec, 72°C 1.5 min), 72°C 7 min, hold at 4°C. Expected band sizes were ∼350 bp for the recombined allele, and ∼1239 bp for the unrecombined/floxed allele. Primers to detect *Ascl1* recombination include the following: Sense *Ascl1* 5’UTR:5’-AAC TTT CCT CCG GGG CTC GTT TC-3’ (for Cre recombined fwd), VR2: 5’-TAG ACG TTG TGG CTG TTG TAG T-3’ (for Cre recombined rev), MF1 5’-CTA CTG TCC AAA CGC AAA GTG G-3’ (for floxed fwd), and VR2 5’-TAG ACG TTG TGG CTG TTG TAG T-3’ (for floxed rev). PCR conditions were 94°C 5 min, 30 cycles of (94°C 1 min, 64°C 1.5 min, 72°C 1 min), 72°C 10 min, hold at 4°C. Expected band sizes were ∼700-850 bp for the *Ascl1* recombined allele, and ∼857 bp for the unrecombined/floxed allele. Recombination PCR to detect *Rbl2/p130* recombined (∼350 bp) and floxed (>1500 bp) alleles was performed under conditions and with primers as previously described^85^. All degrees are in Celsius. PCR products were run on 1.2% agarose/TAE gels containing ethidium bromide or CyberSafe and images were acquired using a BioRad Gel Dox XR imaging system.

##### Generation of basal-organoid derived allografts

After validating recombination in basal organoids (∼3 months after initial CellTagging for CellTagged organoids and ∼1 month post Cre treatment for all organoids), fully-recombined RPM, RPR2, and RPMA basal organoids were implanted as whole or partially digested organoids into flanks of scid/beige mice (Taconic/Charles River) with ∼0.5–3 ×10^6^ cells per flank in 50 μL of 50:50 Matrigel:OCM mix.

Following implantation, basal organoid allografts were measured 1–3x weekly with calipers and were collected when tumours reached an average of 1 cm^3^ but no greater than 2 cm^3^ or upon ulceration, whichever was earlier. Tumour tissue was then subject to FFPE and/or dissociation for single cell RNA-seq (scRNA-seq) experiments and/or re-implantation.

##### CRISPR editing of organoids and validation

To generate *Pten-*knockout RPM and RPMA organoids, basal organoids transformed by CMV-Cre were infected with a non-targeting sgRNA (sg*Ctrl*) or an sgRNA against *Pten* (sg*Pten*: TCA TCA AAG AGA TCG TTA GC) that were cloned into the LCV2-puro backbone (Addgene Plasmid #52961). In brief, high-titer virus (∼1–5 ×10^7^ pfu) was produced using HEK-293T cells transfected with a three-plasmid system including LCV2-sg*Ctrl* or sg*Pten* and lentiviral packaging plasmids pCMV delta R8.2 (Addgene Plasmid #8455) and pCMV-VSVG (Addgene Plasmid #8454). Viruses were harvested at 48 and 72 h post-transfection, concentrated by ultracentrifugation (25,000 RPM for 1.45 hr), resuspended in1X sterile PBS, and stored at -80°C until use. Fully recombined RPM and RPMA organoids were subject to spinoculation with high-titer virus using methods described in “*Lentiviral transduction of organoids with CellTag Library V1”.* Successful editing of *Pten* was validated via T7 endonuclease genome mismatch assays using the following primers: Fwd: 5’-CTCTCGTCGTCTGTCTA-3’; Rev: 5-’ CGAACACTCCCTAGGTGAATAC -3’. Briefly, a ∼1000 bp region containing the sg*Pten* site was amplified using Q5 high-fidelity DNA Polymerase (NEB cat# M0492). PCR conditions used were 98°C 30 sec, 35 cycles of (98°C 10 sec, 65°C 20 sec, 72°C 30 sec), 72°C 2 min, hold at 4°C. PCR product was purified using a Zymo PCR DNA Clean and Concentrator Kit (Zymo cat# D4030), and 200 ng of annealed PCR product was subject to T7 Endonuclease I digestion for 15 min at 37°C. The digestion was quenched with 0.25M EDTA. Products (digested and undigested controls) were run on agarose/TAE gels containing ethidium bromide or CyberSafe and images were acquired using a BioRad Gel Dox XR imaging system.

Immunoblotting was performed to validate PTEN loss via downstream induction of phospho-AKT (Ser473) in RPM and RPMA basal organoids, using methods described above for human cell lines. Primary antibodies included: pAKT (Ser473) (1:1000, CST cat#4060S), total AKT (1:1000, CST cat#9272), and HSP90 (1:1000, CST cat#4877) as loading control.

#### Single cell transcriptomics

##### Preparation of single cell suspensions for scRNA-seq

All organoids were prepared for single-cell transcriptomics ∼3 months after initial CellTaggingand ∼1 month post-Cre treatment (for transformed organoids). Wildtype basal organoids and CMV-Cre-recombined and CellTagged RPM, RPR2, RPMA basal organoids were dissociated to single-cell suspensions using TrypLE Express for 30 min at 37°C, with mechanical disruption every ∼10 min. RPR2 transformed organoids were not analyzed within this study, but data are included in the associated GEO deposition. Allograft and primary tumours used for scRNA-seq were isolated fresh from the lung or flank of mice and immediately subject to digestion and preparation for sequencing. Basal allograft tumours analyzed were: n=1 RPM CMV-Cre; n=4 pooled, multiplexed RPM TAT-Cre; n=3 pooled RPMA CMV-Cre; and n=1 RPR2 CMV-Cre. Primary tumours sequenced for this study included n=2 pooled tumours each from n=2 RPM K5-Cre mice (two tumours from one RPM K5-Cre mouse was multiplexed). Tumour tissue was mechanically dissociated into small clumps using scissors in 1 mL of an enzymatic digestion cocktail per sample then incubated for 30 min at 37°C. The digestion cocktail consisted of 4200 μL of HBSS-free media (Thermo Fisher cat#14175), 600 μL of TrypLE Express (Invitrogen cat#12604013), 600 μL of 10 mg/mL collagenase type 4 (Worthington Biochemical cat# LS004186) prepared in HBSS-free media (Thermo Fisher cat#14175-095), and 600 μL of dispase (50U/mL, Corning cat #354235) and sterilized using a 0.22 μm syringe filter. Enzymatic digestion was quenched on ice with 500 μL of quench media containing 7.2 mL of Leibovitz’s L15 media (Thermo Fisher cat#11415), 800 μL of FBS, and 30 μL of 5 mg/mL DNase (Fisher Sci cat# NC9709009) in HBSS-free media. The tissue suspension was passed through a 100-μm cell strainer. Cells were spun at 2500xg for 5 min at 4°C. Supernatant was removed and replaced with 500 μL of ACK (ammonium-chloride-potassium) lysis buffer per sample to remove red blood cell contamination (3 min incubation at 37°C; VWR cat#10128-808). Reaction was quenched with 10 mL of cold DMEM/F12-Advanced media supplemented with 10% FBS,1% L-glutamine, and 1% Pen/Strep. Cells were spun at 500xg for 5 min at 4°C and resuspended in cold, filtered media or cold, filtered 0.04% BSA in 1X PBS at a concentration of 1–2×10^6 cells/mL and counted manually with a hemocytometer.

##### RNA-seq library preparation

Multiplexed samples included CMV-Cre treated organoids (RPM, RPMA, RPR2) and one K5-Cre RPM sample containing one tumour more central in the lung and one tumour nearer the trachea. Samples were multiplexed before library preparation using 10X Genomics 3’ CellPlex Kit Set A (cat#1000261) and Feature Barcodes (cat#1000262), and following 10X Genomics Demonstrated Protocol (CG000391) following suggestions for “Dissociated Tumor Cells”. After CellPlexing, samples were loaded onto a 10X Chromium X series controller (10X Genomics cat#1000331), targeting 10,000 cells per sample. Samples not subject to multiplexing were immediately loaded onto a 10X Chromium X controller targeting 10,000–20,000 cells per sample. Library preparation was performed following manufacturer’s protocols using the 10X Chromium Next GEM Single Cell 3′ Kit, v3.1 (10X Genomics cat# PN-1000268).

Completed libraries were sequenced on an Illumina NovaSeq 6000, Illumina NextSeq 1000, or a NovaSeq X Plus to target >30,000 reads per cell with the 10X-recommended paired end sequencing mode for dual indexed samples. Individual sample details, including CellPlex oligo information, are provided as metadata with the GEO submission. ScRNA-seq data from n=5 RPM CGRP-Cre initiated primary tumours used in this study were previously published (GEO: GSE149180 and GSE1555692)^42,43^.

##### Demultiplexing and data alignment

ScRNA-seq data were demultiplexed and processed into FASTQ files via the 10X Cell Ranger v7.2.0 pipeline. Primary tumour samples were aligned to a custom mouse genome (GRCm38-mm10-2020-A) including *eGFP*, *Cas9,* firefly *Luciferase* (*fLuc*) and *Venus* to detect recombined alleles in our various mouse models. RPM tumours in this publication express *fLuc*^68^ following recombination of the *MycT58A^LSL/LSL^*allele, and RPMA tumours express *fLuc* and *Venus* following recombination of the *MycT58A^LSL/LSL^* and *Ascl1^fl/fl^* allele, respectively. CellTagged basal allograft tumour samples were aligned to a custom mouse genome (GRCm38-mm10-2020-A) with *fLuc* and *Venus* to aid in detecting recombined tumour cells and include *GFP.CDS* and *CellTag.UTR* transcripts to allow detection of CellTags, according to the published workflow^96,97^ (https://github.com/morris-lab/CellTagR). Sequences used for custom genome builds are included as supplementary files in this publication’s GEO deposit (GSE279200). Count barcodes and UMIs were generated using cellranger count or cellranger multi pipelines for CellPlexed samples.

##### Initial quality control and normalization

Quality control and downstream analysis were performed in Python (v3.8.8) utilizing Scanpy (v1.10.0), according to current expert recommendations for single cell best practices^129^. Anndata objects were created from filtered feature matrices with sc.read_ 10X_mtx(), and quality metrics were calculated using sc.pp.calculate_qc_metrics(). Low quality cells and potential doublets were excluded by selecting for cells with ≤15% mitochondrial content and the following criteria for respective sample types: allograft tumours and organoid samples >1000 total counts, between 500 and 7000 genes with at least one read; RPM primary tumours, >1000 total counts, between 2000 and 10,000 genes with at least one read. Normalized counts were calculated with sc.pp.normalize_total() and a target sum of 10,000. Integrated anndata objects containing multiple scRNA-seq datasets were combined using adata.concatenate() with join=’outer’.

##### Additional quality control and clustering

Data were further processed using Scanpy (v1.10.0) and scVI-tools (v0.17.4) and benchmarking standards to minimize batch effects while maintaining true biological variability, particularly across integrated objects, ^130^. Specifically, a probabilistic deep learning model was developed using scvi.model.SCVI.setup_anndata() to initialize the integration and clustering of datasets from concatenated anndata objects containing all samples and genes, continuous covariate keys set as percent mitochondrial counts, and batch keys identifying samples prepared or sequenced at different times. The model was trained with default parameters, an early stopping patience of 20, and a maximum of 500 epochs, using the model.train() function. The latent representation of the model was obtained with model.get_latent_representation() and added to the .obsm of each anndata object. Neighbors were then calculated with sc.pp.neighbors() with use_rep set to the .obsm category added from the latent representation. UMAP embedding was performed using sc.tl.umap()with min_dist=0.5. Finally, Leiden clusters were generated with sc.tl.leiden() with resolution set to 1.0 for initial steps. As is required throughout the scVI pipeline, we utilized raw counts for all steps above.

Additional rounds of quality control and data filtering were performed per dataset by assessing n_genes_by_counts, total counts, and percent mitochondrial counts per cluster. In general, the model tends to cluster low quality and doublet cells together, so clusters with exceptionally high or low average genes_by_counts, total counts, or mitochondrial content were labeled low-quality and considered for removal from the dataset. Removal was performed after assessing gene expression based on known markers of tumour and normal lung cells, and marker genes per cluster as determined by sc.tl_rank_genes_groups(), to help ensure that biological cells that normally have higher or lower n_genes_by_counts, total_counts, or percent mitochondrial content were not aberrantly filtered out. In addition, in tumour samples from autochthonous or allograft models, we removed non-tumour cells by assessing common immune and normal lung cell type gene expression. Each time a cluster was removed, we ran the scVI pipeline on the new anndata object iteratively through this quality control step until there were no longer any low-quality cell clusters in the anndata object. Additionally, each time clusters were removed from a larger anndata object, the pipeline was re-run for optimal clustering. Final Leiden clusters were determined with sc.tl.leiden() resolution set to 0.5 for all datasets except primary RPM tumours, which was set to 0.25 based on heterogeneity in marker expression determined to be biologically relevant. Source code to reproduce analyses methods is on GitHub (https://github.com/TOliver-Lab/Ireland_Basal_SCLC_2024) and publicly accessible upon publication.

##### Plot generation

UMAP plots showing clustering, sample information, and/or expression of specific genes were generated with sc.pl.umap() using normalized counts as input. Dotplots of normalized counts were generated with sc.pl.dotplot() and, if clustered, dendrogram set to True. For violin plots, data were first converted from anndata objects to Seurat objects using the readH5AD() function in the zellkonverter package and the CreateSeuratObject() function in the SeuratObject package then plotted in R using Seurat’s VlnPlot() function or the plotColData function in the scater package.

##### Transcriptomic gene signatures and differential gene expression analysis

For signature score assessment, and differential gene expression analysis, anndata objects were converted to Seurat objects in R and normalized and log transformed counts were used for visualization after the raw count data were subject to Seurat’s NormalizeData() function. Cell cycle scores were assigned to Seurat objects using the CellCycleScoring() function and Seurat’s cc.genes gene lists, converted to mouse homologs. MYC^42^, ASCL1^82^, and NEUROD1^43^ target gene signatures were previously described, and scores for each gene signature were added to metadata of converted Seurat objects using the AddModuleScore() function. Each target gene score represents conserved transcriptional targets identified via ChIP-seq +/-RNA-seq on mouse and human SCLC cell lines and/or tumours. To generate a POU2F3 target gene signature, published .bed files from POU2F3 ChIP on two human POU2F3+ SCLC cell lines, H526 and H1048, were downloaded (GEO: GSE247951)^31^. Peaks were called and annotated using ChIPseeker in R. Genes with peaks in the promoter region (<2kb from gene TSS) were considered target genes. Conserved target genes between H1048 and H526 were identified, and then only conserved target genes also enriched by log2FC >0.5 in POU2F3-high versus -low human SCLC tumours by published bulk RNA-seq^38^ were included in the final POU2F3 target gene score, which was added to Seurat metadata as described above. ASCL1, NEUROD1, POU2F3, and MYC target gene scores are included in Supplementary Table 2. Normal NE, Tuft, and Basal cell scores were previously published as consensus gene lists derived from mouse scRNA-seq data on normal lung cell types^14^, but proliferation genes, which included ‘Rpl-‘ and ‘Rps-‘ genes were removed to eliminate cell cycle differences and focus on fate-specific markers. The published Ionocyte consensus signature was limited to only 19 genes^14^, so we applied an expanded Ionocyte signature derived from genes >0.5 log2FC enriched in droplet-based scRNA-seq data on mouse trachea/lungs from the same study^14^ (63 genes total). In addition, we utilized a human Ionocyte signature derived from the top 100 human ionocyte markers established in the same study from scRNA-seq on human bronchial epithelium^14^. Resulting normal cell type gene signatures are in Supplementary Table 2. SCLC archetype signatures (A, N, and P) are previously published^46^ and derived from archetype assignments of human SCLC cell lines. Archetype signatures per subtype are in Supplementary Table 2. Gene sets for human SCLC subtypes A, N, and P (Extended Data Fig. 2d) were obtained from scRNA-seq data on human tumours and are published^81^. Inflamed SCLC tumour signatures are derived from published non-negative matrix factorization (NMF) studies on bulk RNA-seq data from n=81 human SCLC tumours^12,65^ (Gay et al) or mRNA, protein, and phosphorylation data from n=107 human SCLC tumours^38^ (Liu et al), where distinct inflammatory subsets were identified (annotated as NMF3 in both studies). The Gay et al Inflammatory signature (Fig. 6f) comprised genes >1 log2FC enriched versus other SCLC subtypes from the NMF-derived gene signature (n=1300 total genes)^12^, resulting in a signature with n=379 human genes converted to mouse homologs (Supplementary Table 6). The Liu et al Inflammatory signature comprised the top 100 genes enriched in NMF3 vs other NMF groups (>1 log2FC)^38^, converted to mouse homologs (Supplementary Table 6). Lastly, NE score was determined based on Spearman correlation with an established expression vector, where ∼41 NE and ∼87 non-NE human cell lines were used to identify a core 50-gene signature comprised of 25 NE genes and 25 non-NE genes that robustly predict NE phenotype^83^. Seurat objects were converted to Single Cell Experiment^131^ objects with the as.SingleCellExperiment() function, then NE score was added as metadata and visualized using the Scater plotColTable() function. Other signatures were visualized similarly with Scanpy, or using FeaturePlot() and/or VlnPlot() functions in Seurat. Marker genes of Leiden clusters in UMAP-embedded space of Figs. 1i, 2e, 2m, and Extended Data Fig. 4a (Supplementary Table 1) and UMAP of Fig. 3f/FA map of Fig. 4c (Supplementary Table 3) were calculated using the sc.tl.rank_genes_groups() on normalized and log-transformed counts with Wilcoxon rank-sum and number of genes set to 500.

##### CellTag analysis and clone calling

CellTags were identified in the RPM and RPMA basal organoid and allograft scRNA-seq samples using processed BAM files (from 10X CellRanger count output, see above methods), and following the CellTagR pipeline documentation (https://github.com/morris-lab/CellTagR). In short, BAM files were filtered to exclude unmapped reads and include reads that align to the *GFP.CDS* transgene or *CellTag.UTR*. CellTag objects were created in R, and CellTags were extracted from the filtered BAM files to generate matrices of cell barcodes, 10X UMIs, and CellTags. The matrix was further filtered to include only barcodes identified as cells by the CellRanger pipeline, and then subject to error correction via Starcode. CellTags not detected in our Whitelist with representation in the >75^th^ percentile (generated from assessment of our lentiviral library complexity, described above) were also removed. Clones were assigned as cells expressing >2 but <20 CellTags with similar combinations of CellTags (Jaccard similarity better than 0.8). For scRNA-seq analysis and CellTag visualization, we used Scanpy (v1.10.0) for initial QC and scvi-tools (v0.17.4) for integration and clustering in Python, following methods described above. Clonal representation pre-and post-implant (Extended Data Fig. 6a) were visualized in R using ggalluvial plots, as a percentage of all clones captured per sample (all clones >2 cells included). Clonal analyses beyond this include only clones identified in RPMA and RPM allograft tumour samples with >10 cells per clone (>5 cells per clone upon stringent QC) (n=12 RPM, n=40 RPMA), to ensure robust sampling of each clone. CellTag-based clonal information was added as metadata by 10X-assigned barcode to visualize clone distribution and cell identity per clone using standard visualization functions in R. CellTag metadata for RPM and RPMA allograft tumour samples are included in Supplementary Table 4. CellTagging on RPR2 basal organoids and allografts occurred, but clonal representation was limited in the allograft tumour due to long latency of this model (∼6 mo) and strong bottlenecks; thus, no CellTag information on RPR2 samples are included in this study.

##### FA mapping, Diffusion Pseudotime, and CellRank analyses

To visualize clonal trajectories, scRNA-seq data from RPM and RPMA basal-derived allografts were projected via force-directed graphing with ForceAtlas2^98^ in Scanpy using the sc.pl.draw_graph() function with default settings. Force-atlas mapping revealed ∼14 cells of Leiden cluster 10 as outliers that were removed prior to diffusion pseudotime computation to allow for a more continuous predicted pseudotime trajectory. We chose to model combined RPM and RPMA allograft cells on one trajectory, because we observed that cells from each genotype occupied all Leiden clusters, albeit at variable frequencies, suggesting cells in each model have the potential to reach all transcriptional phenotypes. After removal of outlier cells, sc.tl.draw_graph was re-run on the subsetted object and cells were projected in FA space.

Diffusion pseudotime was calculated with sc.tl.dpt() with default settings (n_dcs=10) after setting root cells as Cluster 10 basal-like cells (the phenotypic starting point of the experiment, i.e. basal organoids) with adata.uns[‘iroot’]. Next, diffusion pseudotime was used as input to perform CellRank2 analysis^132^, which computes a transition matrix of cellular dynamics and uses estimators to calculate subsequent fate probabilities, driver genes, and gene expression trends. First, we set a PseudotimeKernel (pk), then calculated a cell-cell transition matrix using pk.compute_transition_matrix(). Next, the Generalized Perron Cluster-Cluster Analysis (GPCCA) estimator, a Markov chain based estimator in CellRank^133^, was initialized with g=GPCCA(pk). To assign macrostates, the estimator was fit using g.fit() with n_states set to 11 and cluster_key set to assigned Leiden clusters. Setting n_states to 11 allowed for all Leiden clusters with distinct SCLC fate phenotypes to be picked up as macrostates by the estimator. Cluster 10, the basal-like cluster, was assigned to the estimator as an initial state using g.set_initial_states(), consistent with the assigned pseudotime root state. Terminal states were predicted by the estimator based on highest stability values, using default settings with the g.predict_terminal_states() function and allow_overlap set to True. Predicted terminal states are depicted in Extended Data Fig. 6e. Next, g.compute_fate_probabilities() was used to assign probability values to all cells based on how likely they each are to reach each terminal state. Fate probabilities per cell, annotated by Leiden cluster, were visualized using cr.pl.circular_projection(). The resulting graph was additionally manually annotated with assigned cell fate groupings for Fig. 4i. Predicted lineage drivers of each terminal state were computed with the g.compute_lineage_drivers() function that predicts driver genes by correlating gene expression with fate probability values. Predicted lineage drivers for each terminal state are available in Supplementary Table 4. Finally, we focused on individual trajectories of interest to visualize temporal expression patterns of predicted driver genes along pseudotime using Generalized Additive Models (GAMs). To initialize a model for GAM fitting, the cr.models.GAM() function was run with default parameters and max_iter set to 6000. The top 50 predicted driver genes (or top 200 for the hybrid NE/Neuronal, Cluster 2 trajectory) were fit to the GAM model, to successfully fit ∼50 total genes per trajectory of interest. Gene expression changes over pseudotime were visualized using the cr.pl.heatmap() function and sorted according to peak expression in pseudotime (Fig. 4j). Key parameters for the cr.pl.heatmap() function included data_key set to normalized counts, show_fate_probabilities set to True, time_key set to diffusion pseudotime, and show_all_genes set to True. Source code to reproduce these analyses are deposited at Github (https://github.com/TOliver-Lab/Ireland_Basal_SCLC_2024).

#### Human SCLC data from Caris Life Sciences

##### Whole Transcriptome Sequencing (WTS) sample preparation and data alignment

WTS uses a hybrid-capture method to pull down the full transcriptome from formalin-fixed paraffin-embedded (FFPE) tumour samples using the Agilent SureSelect Human All Exon V7 bait panel (Agilent Technologies, Santa Clara, CA) and the Illumina NovaSeq platform (Illumina, Inc., San Diego, CA).

FFPE specimens underwent pathology review to discern the percent tumour content and tumour size; a minimum of 10% tumour content in the area selected for microdissection was required to enable enrichment and extraction of tumour-specific RNA. Qiagen RNA FFPE tissue extraction kit was used for extraction, and the RNA quality and quantity were determined using the Agilent TapeStation. Biotinylated RNA baits were hybridized to the synthesized and purified cDNA targets, and the bait-target complexes were amplified in a post-capture PCR reaction. The resultant libraries were quantified and normalized, and the pooled libraries were denatured, diluted, and sequenced. Raw data was demultiplexed using the Illumina DRAGEN FFPE accelerator. FASTQ files were aligned with STAR aligner (Alex Dobin, release 2.7.4a Github). Expression data was produced using Salmon, which provides fast and bias-aware quantification of transcript expression^134^. BAM files from STAR aligner were further processed for RNA variants using a proprietary custom detection pipeline. The reference genome used was GRCh37/hg19, and analytical validation of this test demonstrated ≥97% Positive Percent Agreement (PPA), ≥99% Negative Percent Agreement (NPA), and ≥99% Overall Percent Agreement (OPA) with a validated comparator method.

##### Gene expression profiling and SCLC subtype classification

For stratification of patient samples into subgroups, RNA expression values for established and putative lineage-defining transcription factors *ASCL1* (A), *NEUROD1* (N), *POU2F3* (P), and *YAP1* (Y) were standardized to Z-scores. Samples with a single positive Z-score among A/N/P/Y were assigned to the respective gene-associated subgroups, samples with multiple positive Z-scores were classified as ‘Mixed’, and samples with negative Z-scores for all four genes were classified as ‘Lineage-Negative’ (Lin^-^).

##### Gene set enrichment analyses (GSEA)

GSEA was performed using human homologs of normal tuft, basal, neuroendocrine, and ionocyte cell signatures, previously established from mouse and/or human scRNA-seq datasets^14^, as described in above scRNA-seq-related methods and included in Supplementary Table 2. Input included rank-ordered gene lists for each subtype classification by log-fold change expression (log fold-change of ASCL1 subtype versus all other samples, NEUROD1 subtype versus all other samples, etc.). Seurat’s AddModuleScore() function was used to apply hSCLC-A, -N, -Y, -P, -Mixed, and Lin ^-^ signatures from the real-world tumour data to mouse tumour scRNA-seq data, using mouse homologs of the top 100 enriched genes per human subtype vs all other samples.

### Quantification and statistical analyses

Any remaining statistical analysis was performed using GraphPad Prism. Error bars show mean ± SD unless otherwise specified. Significance was determined using appropriate tests as indicated per figure legend and with 95% confidence intervals and p values <0.05 considered statistically significant, unless otherwise indicated. Additional statistical methods related to quantification of immunohistochemistry, and bioinformatic analyses of scRNA-seq can be found in method details. No statistical methods were used to predetermine sample sizes.

## Acknowledgements

We thank members of the Oliver Lab for technical assistance and mouse colony management and administrative support (C. Cheng). We appreciate MedGenome and Duke High Throughput Genomics Core for sequencing services, Duke Cancer Institute Flow Cytometry core for cell sorting, Duke Human Vaccine Institute for use of the Agilent Tapestation, the Preclinical Research Resource (PRR) Core at Huntsman Cancer Institute at the University of Utah for organoid culture reagents, and Jane Johnson (UTSW) for *Ascl1* conditional mice. We also thank the BioRepository & Precision Pathology Center (BRPC), a shared resource of the Duke University School of Medicine and DCI. The BRPC receives support from the P30 Cancer Center Support Grant (P30 CA014236) and from the Cooperative Human Tissue Network (UM1 CA239755). We acknowledge funding sources including the NIH National Cancer Institute (NCI) for awards R01CA262134 (TGO), U24CA213274 (TGO, CMR), U01CA231844 (TGO, RG), F31 CA275295-01 (ASI), R50 CA243783 (DRT), and R35 CA263816 (CMR). We thank the Duke Cancer Institute (DCI) for pilot funds from the P30 Cancer Center Support Grant NIH CA014236.

## Author contributions

Conceptualization: ASI, TGO Methodology: ASI, JMC, DRT, AE, TGO

Investigation: ASI, SBH, BLW, DX, LYZ, MWB, SLR, AE, TGO

Visualization: ASI, AE, TGO

Resources: RG, AD, JCM, SP, CMR, JMC, AE

Funding acquisition: ASI, RG, CMR, TGO Project administration: TGO

Supervision: TGO

Writing – original draft: ASI, TGO Writing – review & editing: all authors

## Ethics declarations

### Competing interests

TGO has a patent related to SCLC subtypes, a sponsored research agreement with Auron Therapeutics, has consulted for Nuage Therapeutics, serves on the scientific advisory board (SAB) for Lung Cancer Research Foundation, and as a consulting editor for *Cancer Research* and *Genes & Development*. CMR has consulted regarding oncology drug development with AbbVie, Amgen, AstraZeneca, Boehringer Ingelheim, and Jazz, and receives licensing fees for DLL3-directed therapies. He serves on the scientific advisory boards of Auron Therapeutics, DISCO, Earli, and Harpoon Therapeutics.

### Diversity, equity, ethics, and inclusion

We worked to include sex balance in the selection of human and non-human subjects. One or more of the authors of this paper self-identifies as a member of the LGBTQIA+ community. One or more of the authors of this paper self-identifies as an underrepresented ethnic minority in their field of research or within their geographical region.

## Additional information

Correspondence and requests for materials should be addressed to Dr. Trudy G. Oliver. Supplementary information is available for this paper.

## Supplementary table legends

**Supplementary Table 1.** Top 500 marker genes of Leiden clusters in Figs. 1i, 2e, 2m, and Extended Data Fig. 4a.

**Supplementary Table 2.** Related to Figs. 2h,i, 3i-k, and Extended Data Figs. 2c–e, 4f and 4i,j, and 9d. ASCL1, NEUROD1, MYC, and POU2F3 target gene signatures, conserved in mouse and human datasets. Normal NE, Tuft, Basal, and Ionocyte cell signatures derived from mouse or human scRNA-seq and previously published^14^. Human SCLC Archetype gene signatures established previously from bulk RNA-seq data on human SCLC cell lines^46^.

**Supplementary Table 3.** Top 500 marker genes of Leiden clusters in Figs. 3f and 4c.

**Supplementary Table 4.** Related to Figs. 4d and Extended Data Fig. 6a-c. CellTag metadata for RPM and RPMA basal organoids and resulting allograft tumours.

**Supplementary Table 5.** Related to Figs. 4i,j and and Extended Data Fig. 6e,f. Putative lineage drivers per CellRank-predicted lineage trajectory in RPM and RPMA basal-derived allograft tumours, predicted with a Generalized Perron Cluster-Cluster Analysis (GPCCA) estimator. Estimator predicts lineage drivers per trajectory by correlating gene expression changes with calculated fate probability values; thus, higher correlation values are stronger predicted drivers.

**Supplementary Table 6.** Related to Fig. 6d-f. Top 100 marker genes per human SCLC subtype derived from the real-world bulk RNA-seq dataset and used to generate human subtype signatures. Antigen presentation, T-cell inflamed, and SCLC-“Inflamed” signatures.

---

**Extended Data Fig. 1:**
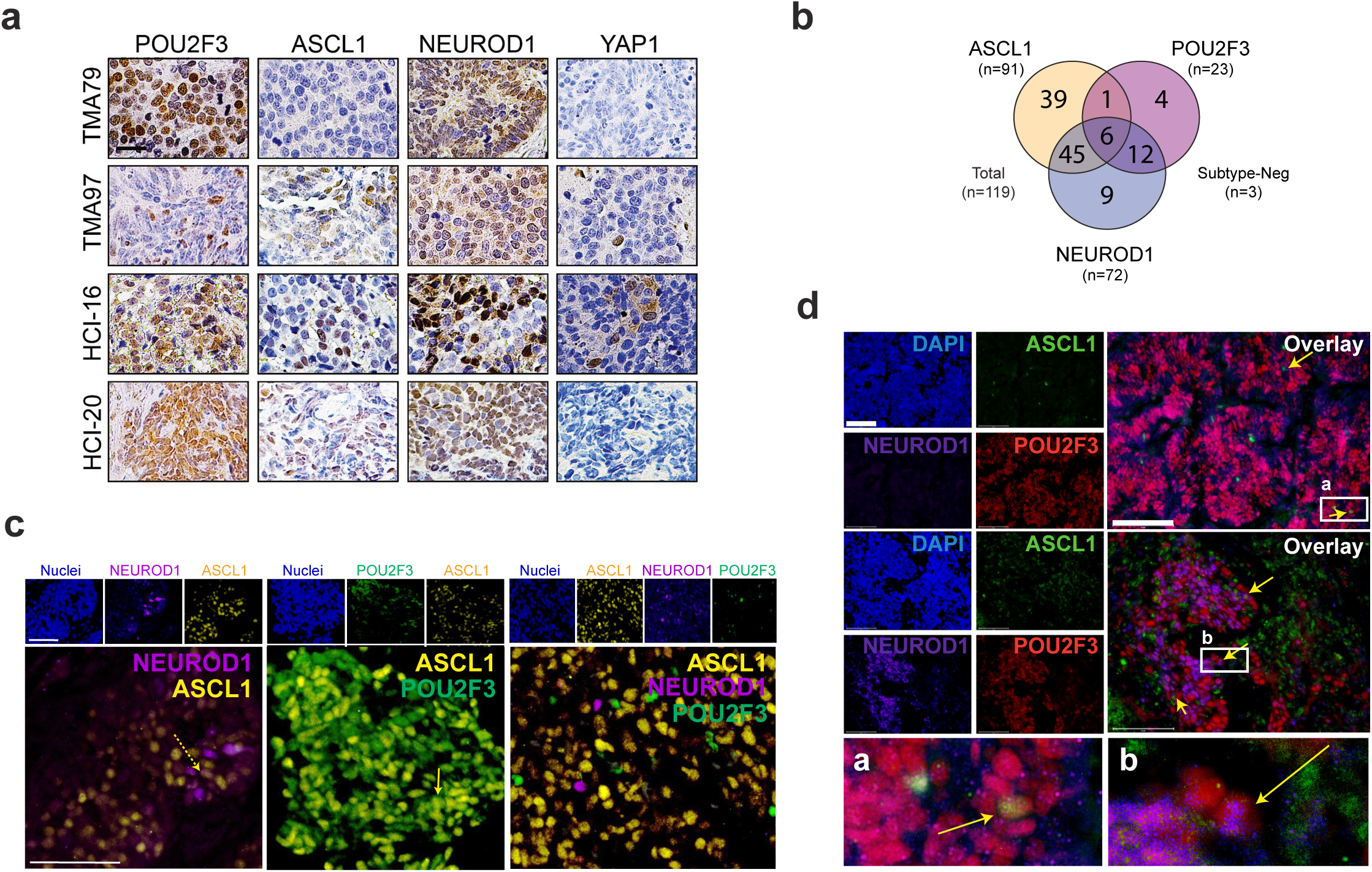
POU2F3^+^ tumours exhibit intratumoural subtype heterogeneity indicative of plasticity. **(a)** Representative images from IHC staining on human SCLC biopsies for given markers. One row = one tumour. Scale bar=25 μm. **(b)** Venn diagram depicting number of human SCLC biopsies (n=119 total) staining positive for ASCL1, NEUROD1, POU2F3, or lacking all markers (“Subtype-Neg”). **(c)** Representative images from co-IF staining on human SCLC biopsies for DAPI (nuclei, blue), ASCL1 (yellow), NEUROD1 (purple) and POU2F3 (green). Individual channels (top) and an overlay without DAPI (bottom) are shown. Scale bars=50 μm (n=28 biopsies stained). **(d)** Representative images from co-IF staining on patient-derived-xenografts (PDX, n=2 distinct models) for DAPI (nuclei, blue), ASCL1 (green), NEUROD1 (purple) and POU2F3 (red). Individual channels (left) and an overlay without DAPI (right) are shown. Yellow arrows and insets (a-b) emphasize co-expressing cells (bottom). Scale bars=75 μm.

**Extended Data Fig. 2:**
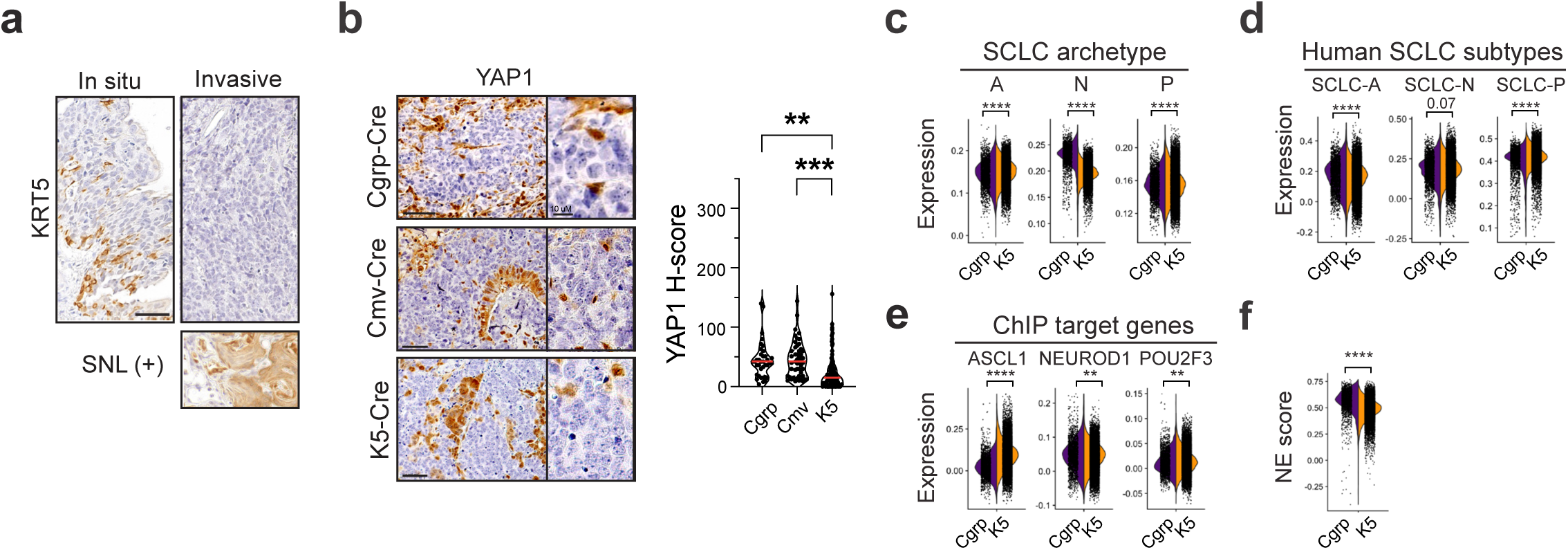
Basal cells give rise to SCLC with expansive subtype heterogeneity. Related to Fig. 1 and Supplementary Table 2. **(a)** Representative IHC images from RPM K5-Cre tumours for KRT5. A *Sox2^LSL/LSL^;Nkx2-1^fl/fl^;Lkb1^fl/fl^* mouse lung squamous tumour^128^ is included as a positive control. **(b)** Representative IHC for YAP1 in RPM tumours initiated by indicated Ad-Cre viruses (left) and corresponding H-score quantification (right). Median (red bar) and upper and lower quartiles (dotted lines) indicated. One-way ANOVA with Tukey’s correction (e,f). *** p<0.0003, ** p<0.003. **(c)** Split violin plot depicting expression of SCLC archetype signatures^46^ per tumour cell by scRNA-seq in CGRP (purple, left) and K5 (orange, right) tumours in Fig. 1h. **(d)** Split violin plot depicting transcriptional signatures of human ASCL1, NEUROD1, or POU2F3^+^ tumours derived from scRNA-seq data^81^ or **(e)** expression of ASCL1, NEUROD1, and POU2F3 target gene signatures derived from ChIP-seq data^31,42,43,113^(Supplementary Table 2) by scRNA-seq for all cells in CGRP (purple, left) and K5 (orange, right) tumours. Each dot is one cell. **(f)** Split violin plot depicting NE score^83^ per tumour cell by scRNA-seq in CGRP (purple, left) and K5 (orange, right) tumours. All scale bars=50 μm. Unless otherwise noted, statistical tests are Student’s unpaired t-tests. **** p<0.0001, ** p<0.01, ns=not significant, p>0.05.

**Extended Data Fig. 3:**
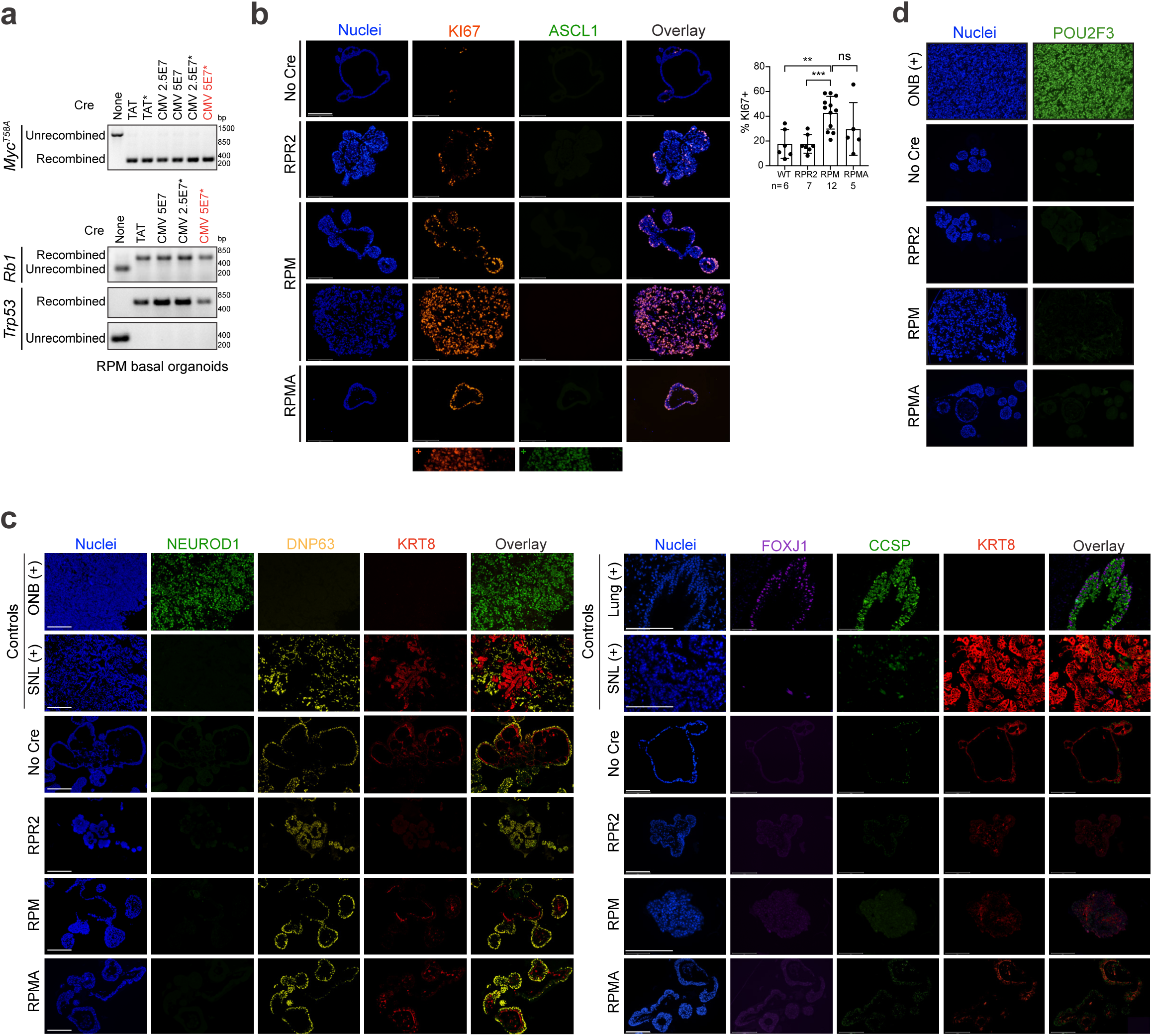
Basal-derived GEMM organoids maintain basal identity in vitro. Related to Figs. 2 and 3. **(a)** Recombination PCR for RPM basal organoids for indicated alleles pre- and post-treatment with TAT-Cre recombinase or Ad-CMV-Cre at two concentrations (2.5e7 or 5e7 pfu). *Organoids subject to spinoculation with CMV-Cre virus. Red font indicates condition used for subsequent allografting. **(b)** Co-IF on basal organoids pre-treatment with Cre (No Cre) or following recombination (RPM, RPR2, RPMA) for DAPI (nuclei, blue), KI67 (proliferation, orange), and ASCL1 (NE cell, green). Positive controls for KI67 and ASCL1 (bottom panel) is an RPR2 SCLC lung tumour. Quantification via CellProfiler^135^ of KI67 positivity per organoid from indicated conditions (right). Number of organoids quantified per group is labeled. One-way ANOVA with Tukey’s correction. *** p<0.003, ** p<0.005, ns=not significant, p>0.05. Error bars represent mean ± SD. **(c)** Co-IF on basal organoids pre-treatment with Cre (No Cre) or following recombination (RPM, RPR2, RPMA): Left) DAPI (nuclei, blue), NEUROD1 (neuronal cell, green), DNP63 (basal cell, yellow) and KRT8 (luminal basal cell, red). Positive control (+) for NEUROD1 is an RPM olfactory neuroblastoma tumour^6^ and for basal markers is an SNL GEMM lung tumour^128^. Right) DAPI (nuclei, blue), FOXJ1 (ciliated cell, purple), CCSP (SCGB1A1, club cell, green) and KRT8 (luminal basal cell, red). Positive control (+) control for FOXJ1 and CCSP is airway from a normal mouse lung, and for KRT8 is an SNL GEMM lung tumour. **(d)** Co-IF on basal organoids pre-treatment with Cre (No Cre) or following recombination (RPM, RPR2, RPMA) for DAPI (nuclei, blue) and POU2F3 (tuft cell, green). Positive control (+) is an RPMA olfactory neuroblastoma tumour^6^. All co-IF scale bars=150 μm.

**Extended Data Fig. 4:**
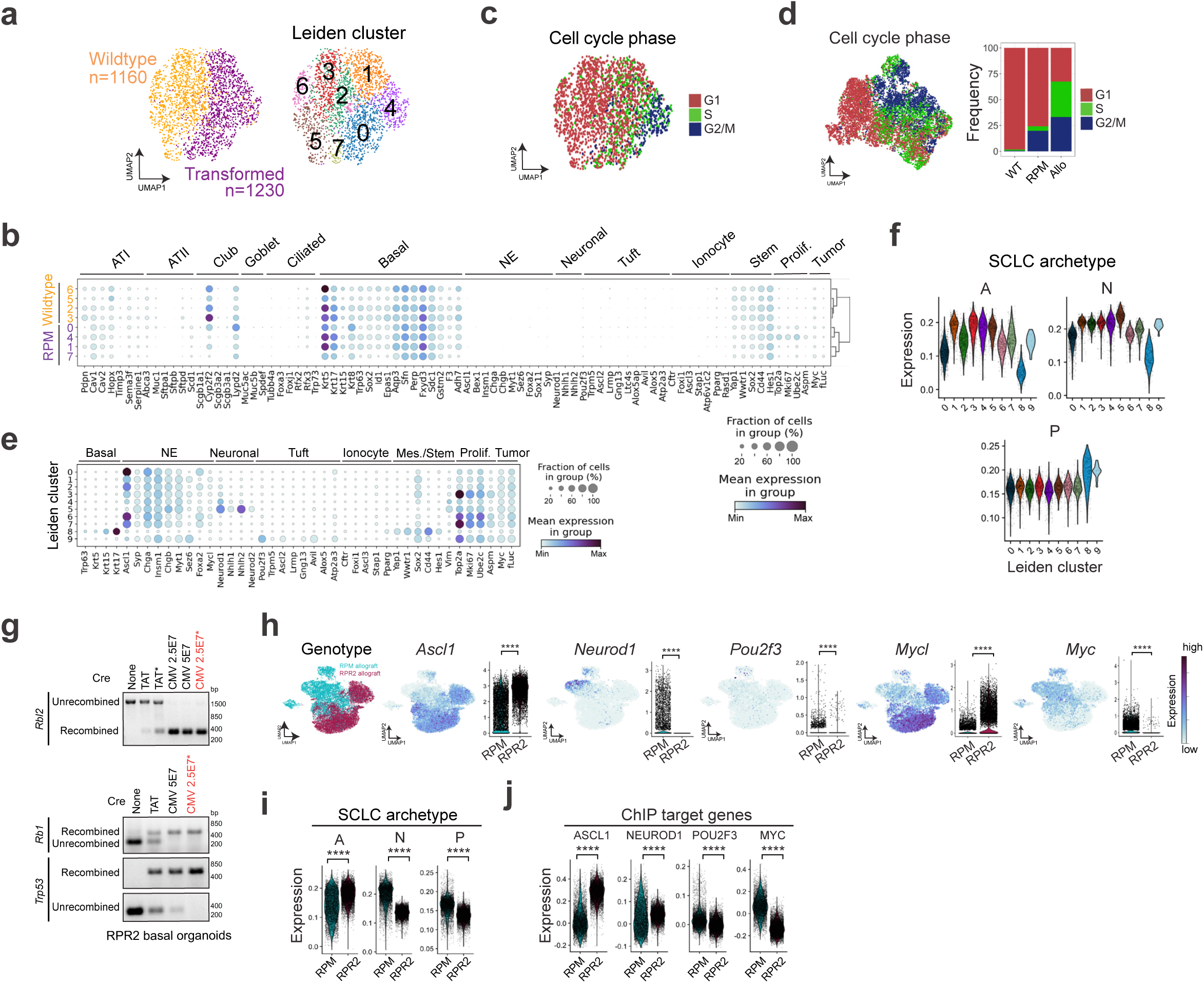
Basal-derived GEMM organoids give rise to neuroendocrine, neuronal, and tuft-like SCLC in vivo. Related to Fig. 2 and Supplementary Tables 1 and 2. **(a)** UMAP of scRNA-seq data from wildtype (orange) and transformed (purple) RPM basal organoids (left) and annotated by Leiden cluster (right) (Supplementary Table 1). **(b)** Dot plot expression of genes marking major lung cell types, stem-like, proliferative, and tumour cells for wildtype versus transformed RPM organoids, grouped by Leiden cluster as assigned in (a). Colour indicates level of gene expression and dot size represents frequency of expression per cluster. **(c)** UMAP of RPM organoids pre- and post-CMV-Cre coloured by cell cycle phase. **(d)** UMAP as in Fig. 2d of RPM organoids pre- and post-CMV-Cre and RPM basal allograft tumour coloured by cell cycle phase (left). Proportion of cells from WT and transformed (RPM) organoid samples and RPM allograft tumour in each phase, represented as % of all cells per sample (right). **(e)** Dot plot expression of genes marking indicated cell types, stem-like, proliferative, and tumour cells for RPM basal allograft tumour by Leiden cluster derived from scRNA-seq in Fig. 2e. **(f)** Violin plot expression of SCLC archetype signatures in RPM basal allograft tumour cells by scRNA-seq as in Fig. 2e, grouped by Leiden cluster. **(g)** Recombination PCR for RPR2 basal organoids for indicated alleles pre- and post-treatment with TAT-Cre recombinase or Ad-CMV-Cre at two concentrations (2.5e7 or 5e7 pfu). *Organoids subject to spinoculation with CMV-Cre virus. Red font indicates condition used for subsequent allografting. **(h)** UMAP and corresponding violin plots depicting expression of indicated SCLC subtype markers and Myc-family oncogenes in RPM versus RPR2 basal organoid allografts from Fig. 2l. **(i)** Violin plot expression of SCLC subtype archetype signatures^46^ or **(j)** ChIP target genes signatures (Supplementary Table 2) per tumour cell in RPM vs RPR2 basal allograft tumour samples from scRNA-seq data in Fig. 2l. A=ASCL1, N=NEUROD1, and P=POU2F3. Unless otherwise noted, statistical tests are Student’s unpaired t-test. **** p<0.0001.

**Extended Data Fig. 5:**
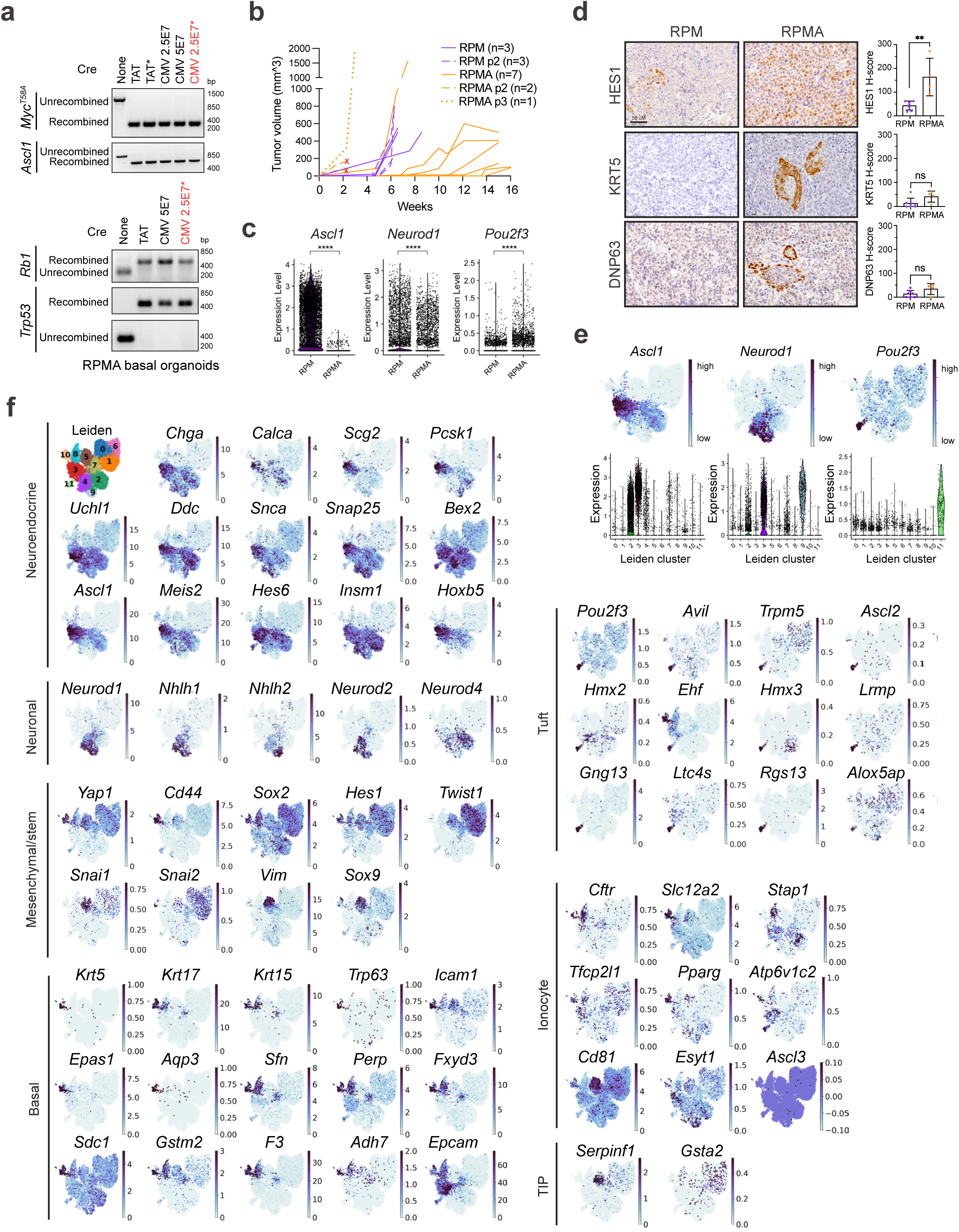
ASCL1 loss promotes POU2F3^+^ tuft-like SCLC. Related to Fig. 3 and Supplementary Table 3. **(a)** Recombination PCR for RPMA basal organoids for indicated alleles pre- (None) and post-treatment with TAT-Cre recombinase or Ad-CMV-Cre at two concentrations (2.5e7 or 5e7 pfu). *Organoids subject to spinoculation with CMV-Cre virus. Red font indicates condition used for subsequent allografting. **(b)** Quantification of tumour volume (mm^3) over time (weeks) in RPM (purple) vs RPMA (orange) basal allografts. Number of tumours quantified are indicated in legend. No suffix=Passage 1 (solid line), p2= Passage 2 (dashed line), p3 = Passage 3 (dotted line). Red “X” indicates censored animals due to early tumour ulceration. **(c)** Violin plot expression of SCLC subtype genes in RPM vs RPMA basal allograft tumours from scRNA-seq data in Fig. 3e. Each dot is one cell. **(d)** Representative IHC images from RPM and RPMA basal-organoid-derived tumours for indicated markers (left). H-score quantification of IHC for indicated proteins (right). Each dot represents one tumour. For each marker, n=3-9 tumours quantified. Scale bars=50 μm. **(e)** FeaturePlots depicting expression of indicated SCLC subtype genes in RPM and RPMA basal allograft tumours from scRNA-seq data in Fig. 3e (top) and corresponding violin plots depicting gene expression per Leiden cluster assigned in Fig. 3f. **(f)** FeaturePlots depicting expression of indicated NE, Neuronal, Mesenchymal/Stem, Tuft, Ionocyte, Tuft-Ionocyte-Progenitor (TIP), and Basal markers in UMAP of RPM and RPMA basal-derived allograft tumours from scRNA-seq data depicted in Fig. 3e. For relevant plots, statistical tests are Student’s unpaired t-test **** p<0.0001, ** p<0.01, ns=not significant, p>0.05. Error bars represent mean ± SD.

**Extended Data Fig. 6:**
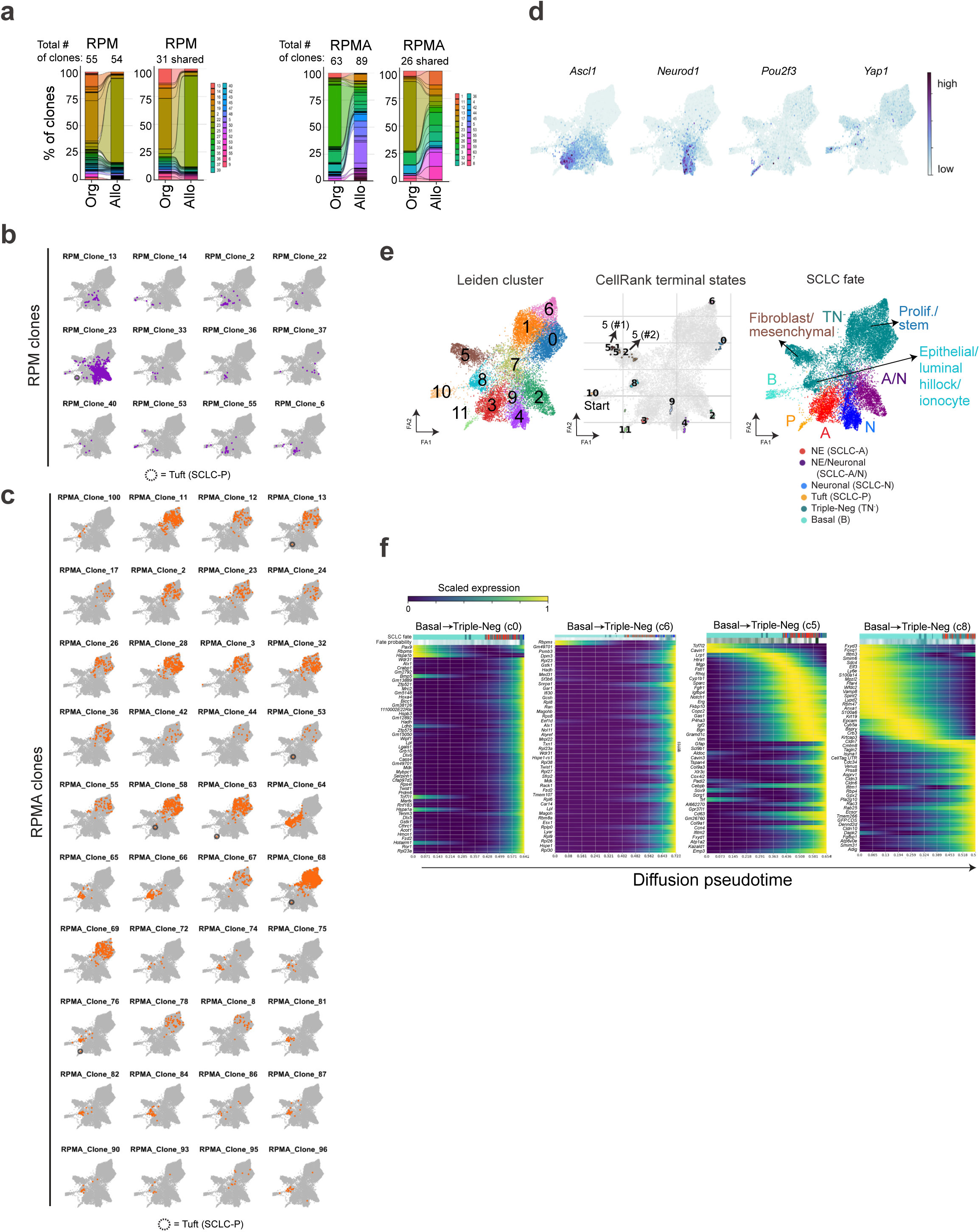
Lineage-tracing reveal distinct SCLC evolutionary trajectories. Related to Fig. 4 and Supplementary Tables 4 and 5. **(a)** Alluvial bar plots depicting number and frequency of CellTag clones captured in RPM and RPMA organoids and resulting allograft tumours. Plots are shown for all clones per genotype (left) as well as only clones shared/captured across both conditions by scRNA-seq (right). Number of clones annotated on figure. **(b)** Projection of individual CellTag clones from RPM basal allograft tumours projected in FA map from Fig. 4c. Dashed circle indicates cells in the Tuft or SCLC-P state in clones. See Supplementary Table 4. **(c)** Projection of individual CellTag clones from RPMA basal allograft tumours projected in FA map from Fig. 4c. Dashed circle indicates cells in the Tuft or SCLC-P state in clones. See Supplementary Table 4. **(d)** UMAP of RPM and RPMA basal allograft tumour cells as in Fig. 4c coloured by expression of SCLC subtype marker genes. **(e)** FA map annotated by Leiden cluster of RPM and RPMA basal allograft tumour cells from Fig. 4c (left) with corresponding CellRank-predicted terminal states (middle) and assigned SCLC cell fate as in Fig. 4c, with added annotations defining phenotypic variation within TN^-^ clusters (right). Predicted CellRank trajectories all had an assigned start of Cluster 10, the cluster most enriched for Basal markers. **(f)** CellRank trajectory-specific expression trends of putative driver genes (Supplementary Table 5), predicted by fitting gene expression and pseudotime coordinates with Generalized Additive Models (GAMs).

**Extended Data Fig. 7:**
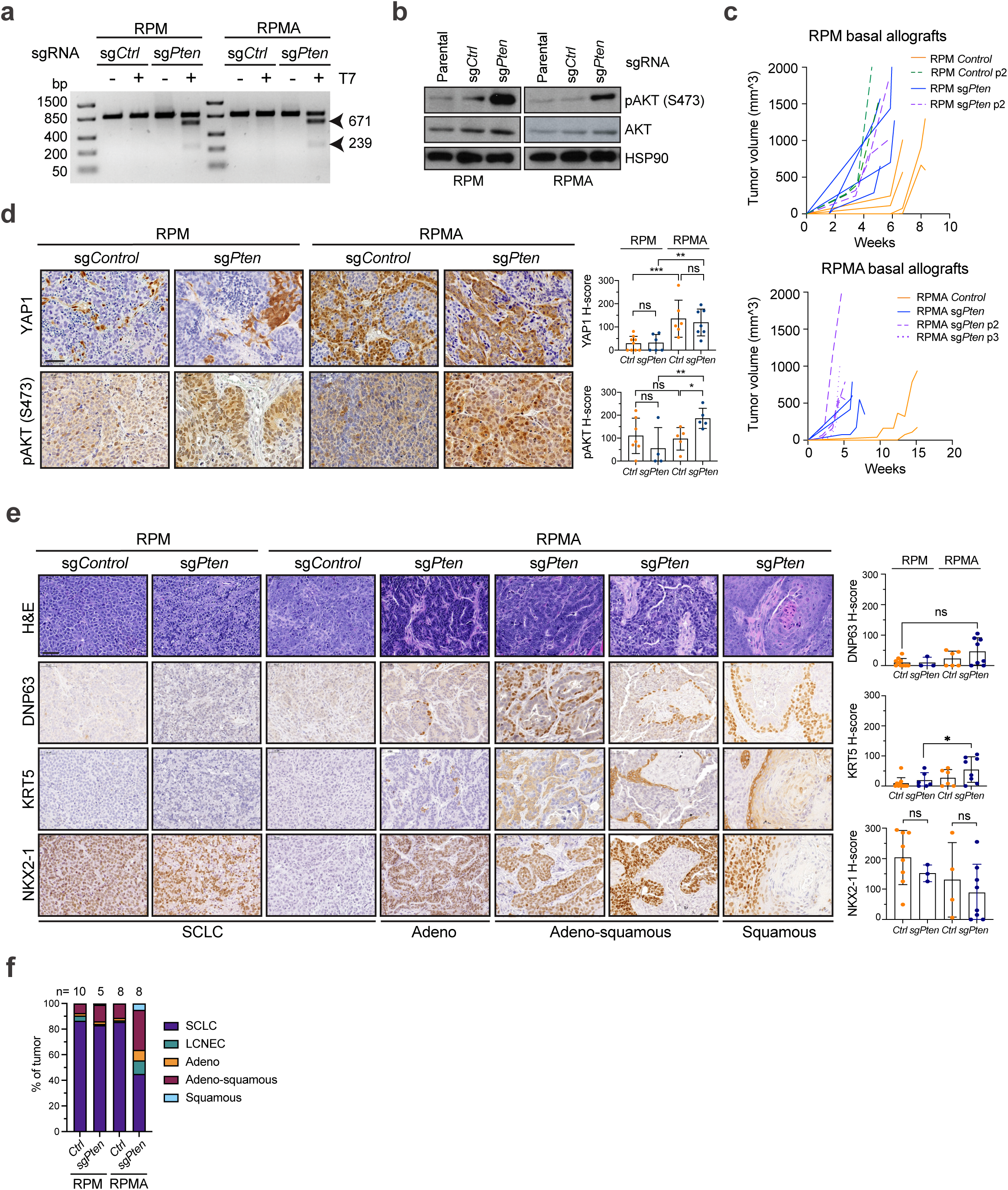
PTEN loss promotes POU2F3 in basal-derived SCLC. Related to Fig. 5. **(a)** T7 endonuclease genome mismatch assay on transformed RPM and RPMA basal organoids after lentiviral infection of LCV2 with non-targeting (sg*Ctrl*) or *Pten-*targeting sgRNAs. Expected products of T7 endonuclease digestion with successful editing are 671 and 239 bp. **(b)** Immunoblot of pAKT (Ser473) and total AKT with HSP90 as a loading control on transformed RPM and RPMA basal organoids after infection with sg*Ctrl* or *Pten-*targeting sgRNAs. **(c)** Quantification of tumour volume (mm^3) over time (weeks) in RPM (left) and RPMA (right) *sgCtrl* (passage 1=orange, solid; passage 2=green, dashed), and *sgPten* (passage 1=blue, solid; passage 2=purple, dashed, passage 3=purple, dotted), basal organoid allografts. **(d)** Representative IHC images from RPM and RPMA basal-organoid-derived parental or LCV2-*sgCtrl* tumours compared to tumours with LCV2-*sgPten* for indicated markers (left). H-score quantification of IHC for indicated proteins. Each dot represents one tumour. For each genotype, n=3-4 parental, n=2-6 sg*Ctrl,* and n=6-9 sg*Pten* tumours were quantified. Parental and sg*Ctrl* tumours are grouped in the same category labeled as *Ctrl.* **(e)** Representative H&E and IHC for diagnostic markers on RPM and RPMA sg*Ctrl* and *sgPten* tumours with SCLC histopathology versus regions of adenocarcinoma (Adeno), adeno-squamous carcinoma (Adeno-squamous), and Squamous carcinoma differentiation. Corresponding H-score quantification on right. For each genotype, n=3-4 parental, n=2-6 sg*Ctrl,* and n=3-8 sg*Pten* tumours were quantified. Parental and sg*Ctrl* tumours are grouped in the same category labeled as *Ctrl.* **(f)** Stacked bar chart depicting average proportions of indicated histopathologies within individual RPM and RPMA control or *Pten*-deleted tumours. Number of tumours analyzed and represented by the average values indicated above stacked bars. Histopathologies determined via analysis of H&E and NKX2-1, P63, KRT5, and SCLC subtype marker staining. LCNEC is large-cell neuroendocrine carcinoma. All scale bars=50 μm. All statistical tests are two-way ANOVA. *** p<0.0008, ** p<0.007, * p<0.04, ns=not significant, p>0.05. Error bars represent mean ± SD.

**Extended Data Fig. 8:**
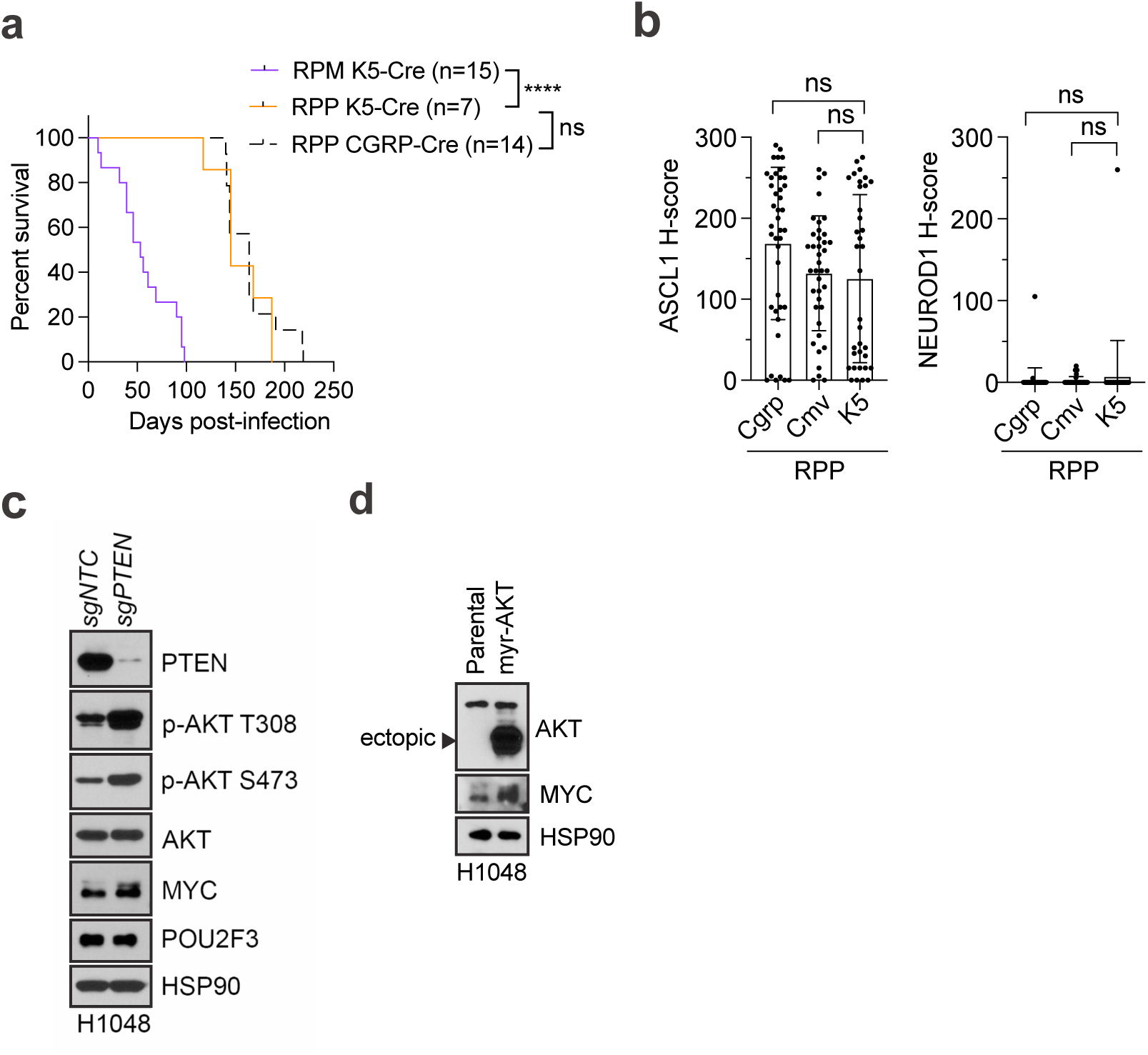
PTEN loss and MYC cooperate to drive POU2F3^+^ SCLC. Related to Fig. 5. **(a)** Survival of RPP mice infected with K5-Cre or Cgrp-Cre compared to RPM mice infected with K5-Cre. Dashed line indicates historical data^68^. Number of mice indicated in the figure. Mantel-Cox log-rank test; **** p<0.0001, ns=not significant, p >0.05. **(b)** H-score quantification of ASCL1 and NEUROD1 in RPP GEMM tumours initiated by indicated Ad-Cre viruses. Each dot is one tumour. N=35-40 tumours from n=5-8 mice per cohort. One-way ANOVA with Tukey’s correction. ns=not significant, p>0.05. **(c)** Immunoblot analysis of human SCLC cell line, H1048, for indicated markers after LentiCRISPRv2 infection with non-targeting (*sgNTC*) or *sgPTEN* sgRNAs. **(d)** Immunoblot analysis of H1048 for indicated markers in parental cells versus cells with ectopic myristoylated-AKT (myrAKT) with HSP90 as a loading control.

**Extended Data Fig. 9:**
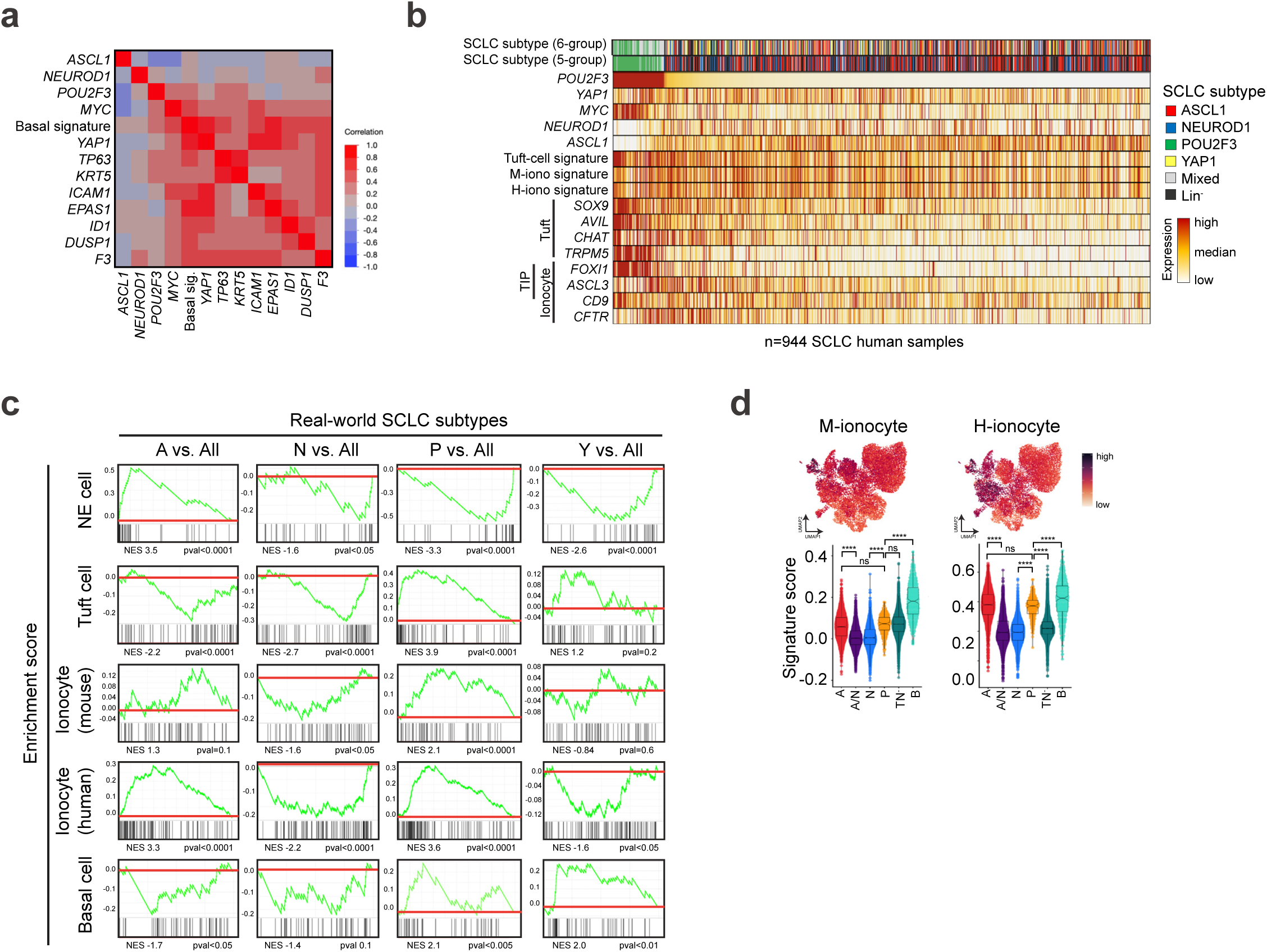
Human SCLC harbours a basal-like subset. Related to Fig. 6 and Supplementary Table 2. **(a)** Spearman correlation matrix depicting individual gene or gene signature correlations by bulk RNA-seq in n=944 human SCLC biopsies. Data include subtype markers, MYC, YAP1, and key basal state markers. **(b)** Heatmap displays expression by bulk RNA-seq of normal tuft, ionocyte, and tuft-ionocyte-progenitor (“TIP”) markers or gene signatures in n=944 human SCLC biopsies, sorted left to right by expression of *POU2F3*. Samples are annotated by classified SCLC subtypes (including the 5-group and 6-group classification schemes). **(c)** Gene set enrichment analysis (GSEA) depicting enrichment of normal neuroendocrine (NE), tuft, basal, and ionocyte cell signatures (Supplementary Table 2) in each human SCLC subtype in the real-world bulk RNA-seq dataset. Normalized enrichment score (NES) and p-values are indicated. **(d)** Expression of ionocyte signatures derived from mouse (M) or human (H) scRNA-seq studies^14^ applied to RPM and RPMA basal allograft tumour cells from Fig. 3e in UMAP space (top) or per assigned SCLC fate in the form of a violin plot (bottom). Box-whisker overlays on violin plots indicate median and upper and lower quartile. One-way ANOVA with Tukey’s correction. **** p<0.0001, ns=not significant, p>0.05.

